# A High-Throughput Screen Reveals the Structure-Activity Relationship of the Antimicrobial Lasso Peptide Ubonodin

**DOI:** 10.1101/2022.12.13.520261

**Authors:** Alina Thokkadam, Truc Do, Xinchun Ran, Mark P. Brynildsen, Zhongyue J. Yang, A. James Link

## Abstract

The *Burkholderia cepacia* complex (Bcc) is a group of bacteria including several opportunistic human pathogens. Immunocompromised individuals and cystic fibrosis patients are especially vulnerable to serious infections by these bacteria, motivating the search for compounds with antimicrobial activity against the Bcc. The natural product ubonodin is a lasso peptide with promising activity against several Bcc species, working by inhibiting RNA polymerase in susceptible bacteria. In this study, we developed a high-throughput screen using next-generation sequencing to examine the fitness of a library of over 90,000 ubonodin variants, generating the most comprehensive dataset on lasso peptide activity so far. This screen revealed information regarding the structure-activity relationship of ubonodin over a large sequence space, indicating certain residues that can tolerate amino acid substitutions and still retain activity. Remarkably, the screen identified one variant with not only improved activity compared to wild-type ubonodin but also a sub-micromolar minimum inhibitory concentration (MIC) against a clinical isolate of the Bcc member *Burkholderia cenocepacia*. Ubonodin and several of the variants identified in this study had a lower MIC against certain Bcc strains than many clinically approved antibiotics. Finally, the large library size enabled us to develop DeepLasso, a deep learning model that can predict the RNAP inhibitory activity of an ubonodin variant.

## Introduction

RiPPs (ribosomally synthesized and post-translationally modified peptides)^1-2^ are a superfamily of natural products defined by their ribosomal origin. RiPP precursor proteins are gene-encoded, translated at the ribosome, and post-translationally modified into their final structure via the action of a suite of enzymes. The gene-encoded nature of RiPPs makes them especially well-suited to high-throughput engineering or structure-activity relationship studies because large libraries of RiPP precursor mutants can be generated using PCR techniques. This ability to generate large libraries of RiPPs has been used to tune existing RiPP functions, such as antimicrobial activity.^3-6^ Using library techniques, RiPPs have also been repurposed to carry out entirely new functions.^7-10^ One family of RiPPs that has been engineered by library-scale mutagenesis is the lasso peptides, named after their unusual threaded structure.^11-12^ This structure comprises an N-terminal macrocyclic ring formed by an isopeptide bond, a loop that threads through the ring forming a mechanical bond, and a C-terminal tail (Figure 1A). Lasso peptides have attracted interest due to their bioactivities, the most well-studied of which is narrow-spectrum antimicrobial activity.^13-22^ A subset of these antimicrobial lasso peptides exerts their function via inhibition of RNA polymerase (RNAP).^23-27^

**Figure 1.**
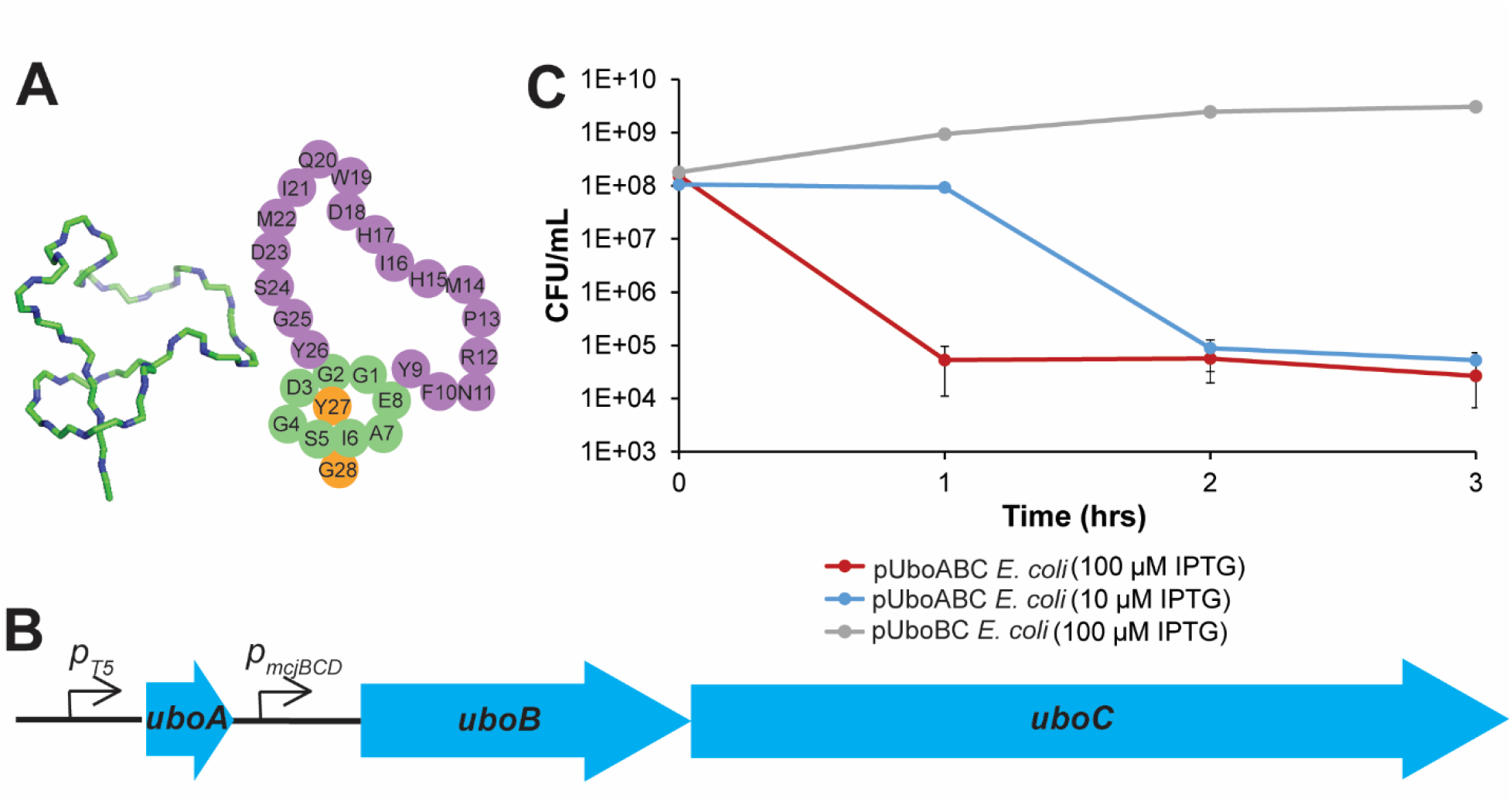
Ubonodin and plasmid constructs. **A**: Structure of ubonodin, drawn from PDB file 6POR, and cartoon representation. **B**: Refactored ubonodin BGC in pUboABC. The *uboA* gene was under the control of an IPTG-inducible T5 promoter. The *uboB* and *uboC* genes were under the control of a constitutive promoter found upstream of the *mcjB, mcjC*, and *mcjD* genes in the microcin J25 BGC. **C**: Viability of cultures of *E. coli* pUboABC and *E. coli* pUboBC. IPTG induction occurred at the 0-hour timepoint. IPTG induction in *E. coli* pUboABC led to a 3-log reduction in the titer of viable cells whereas the *E. coli* pUboBC cells continued to grow normally after induction.

We recently reported the structure and antimicrobial activity of the RNAP-inhibiting lasso peptide ubonodin.^28^ In addition to its unusually large size at 28 amino acids (aa), ubonodin has compelling antimicrobial activity against multiple members of the *Burkholderia cepacia* complex (Bcc).^29^ Bcc members are causative agents of potentially fatal lung disease in people with cystic fibrosis.^30^ Part of our motivation for studying ubonodin is the intrinsic antibiotic resistance in Bcc members^31-35^ underscoring the need for new antimicrobials targeting the Bcc. We have also recently uncovered the molecular basis for the specificity of ubonodin toward Bcc strains using a phenotype-guided approach.^36^ Ubonodin breaches the outer membrane of susceptible Bcc strains by interaction with a specific receptor PupB and crosses the inner membrane by interacting with the ABC transporter YddA. Here we build libraries of all possible single aa variants of ubonodin and a large subset of possible double aa variants of ubonodin. The fitness of each member of these libraries was assessed using a growth-based screen and next-generation sequencing (NGS). This large set of sequence-activity data was used to train a deep learning model, allowing for reliable prediction of activity from the sequence alone. We validated a small subset of the active ubonodin variants against *Burkholderia cenocepacia* clinical isolates using a broth microdilution assay, and even found one variant, ubonodin H17G, that was more potent than wild-type ubonodin. Overall, the results of these screens have given us an unprecedented view into the structure-activity relationship for ubonodin and have identified thousands of ubonodin single and double aa variants with RNAP inhibition activity.

## Results

### Development of a high-throughput screen for RNA polymerase inhibition

The ubonodin biosynthetic gene cluster (BGC) consists of the *uboA, uboB, uboC*, and *uboD* genes. UboA is the precursor to ubonodin while UboB and UboC are a cysteine protease and a lasso cyclase, respectively, and are the enzymes necessary for synthesizing the lasso peptide. UboD is an ABC transporter that pumps ubonodin outside of the cell thus functioning as an immunity factor. We envisioned a screen in which a library of cells, each producing a single ubonodin variant from a plasmid in the cytoplasm, would be grown up *en masse*. Upon induction, cells harboring ubonodin variants capable of inhibiting RNAP would die out while those with either inactive or unprocessed ubonodin variants would survive. By sequencing the plasmids from the cell library pre- and post-ubonodin induction, we would identify which ubonodin variants retain or lose RNAP inhibition activity. To develop such a screen, we constructed the plasmid pUboABC (Figure 1B). This plasmid contains *uboA* under an IPTG-inducible promoter, and the *uboBC* operon under a constitutive promoter. Since the *uboD* gene is not present in this plasmid, induction of wild-type *uboA* is expected to cause cell death due to RNAP inhibition from ubonodin accumulating inside the cell. The plasmid pUboBC, lacking the *uboA* gene, was also constructed as a negative control. Although ubonodin targets *Burkholderia*, the *Burkholderia* and *E. coli* RNAP are highly similar, especially in the regions of the *β* and *β*’ subunits targeted by lasso peptides (Figure S1),^26^ and we have shown previously that ubonodin inhibits *E. coli* RNAP *in vitro*.^28^ As a first test, *E. coli* harboring pUboABC were streaked out on plates with either IPTG to induce *uboA* expression or glucose to repress *uboA* expression. As expected, the pUboABC strain grew on glucose but not IPTG (Figure S2A-B). Next, a spot dilution assay comparing *E. coli* harboring pUboABC or pUboBC indicated a clear growth defect of *E. coli* pUboABC (Figure S2C). The viability of these cultures was also quantitatively determined at various timepoints by measuring colony forming units (CFU) per mL of culture (Figure 1C). The cultures of *E. coli* pUboBC had no growth defect after IPTG induction. However, IPTG induction led to a 3-log reduction in viable cell concentration of *E. coli* pUboABC under induction. Additionally, examination of induced *E. coli* pUboABC cells via brightfield microscopy showed a severe filamentation phenotype (Figure S3) consistent with what has been previously reported for MccJ25.^16^

As a further test of whether these plasmids would be suitable for a library screen, we set up co-culture experiments with *E. coli* pUboABC and *E. coli* pUboBC. The pUboABC and pUboBC plasmids were mixed in a ratio of 9:1 and transformed into *E. coli*, mimicking a library screening workflow. The co-culture was grown up in liquid media and induced at mid-log phase. Samples of the culture were plated at various timepoints, and PCR amplification of the resulting colonies was used to assess the ratio of pUboABC to pUboBC throughout the mock screen. Although pUboBC was present at a low frequency at the beginning of the co-culture, it increased in frequency over time (Figure S4). This plasmid completely overtook the co-culture one hour after induction with 100 μM IPTG and two hours after induction with 10 μM IPTG. This experiment established that cells harboring either an inactive or otherwise non-toxic variant of ubonodin will increase in frequency within the co-culture over time, thus validating this approach for the screening of ubonodin variant libraries.

### Single Mutant Library Construction and Screening

Site saturation mutagenesis was used to construct a library of single mutants of ubonodin. All positions of ubonodin were mutated except the Gly1 and Glu8 residues, which are required to form the isopeptide bond that creates the lasso peptide’s ring. Sites were individually mutated with primers containing NNK codons (where N represents any of the four bases and K represents G or T). NNK codons cover all 20 amino acids with reduced bias relative to the standard genetic code and eliminate 2 of the 3 stop codons. The final plasmid mixtures from mutating each residue were combined in equimolar amounts (Figure S5).

The single mutant library was analyzed by NGS at multiple stages of sample preparation (Figure 2A) to track its evolution throughout the screen. Specifically, the frequency of each possible aa variant was computed at each stage by summing up counts for each distinct translated sequence and dividing by the total number of counts for all variants (Equation S1, SI Methods). Since cells expressing a RNAP-inhibiting ubonodin variant will have growth defects relative to cells with a non-toxic variant, RNAP-inhibiting ubonodin variants are expected to decrease in frequency throughout the screen.

**Figure 2.**
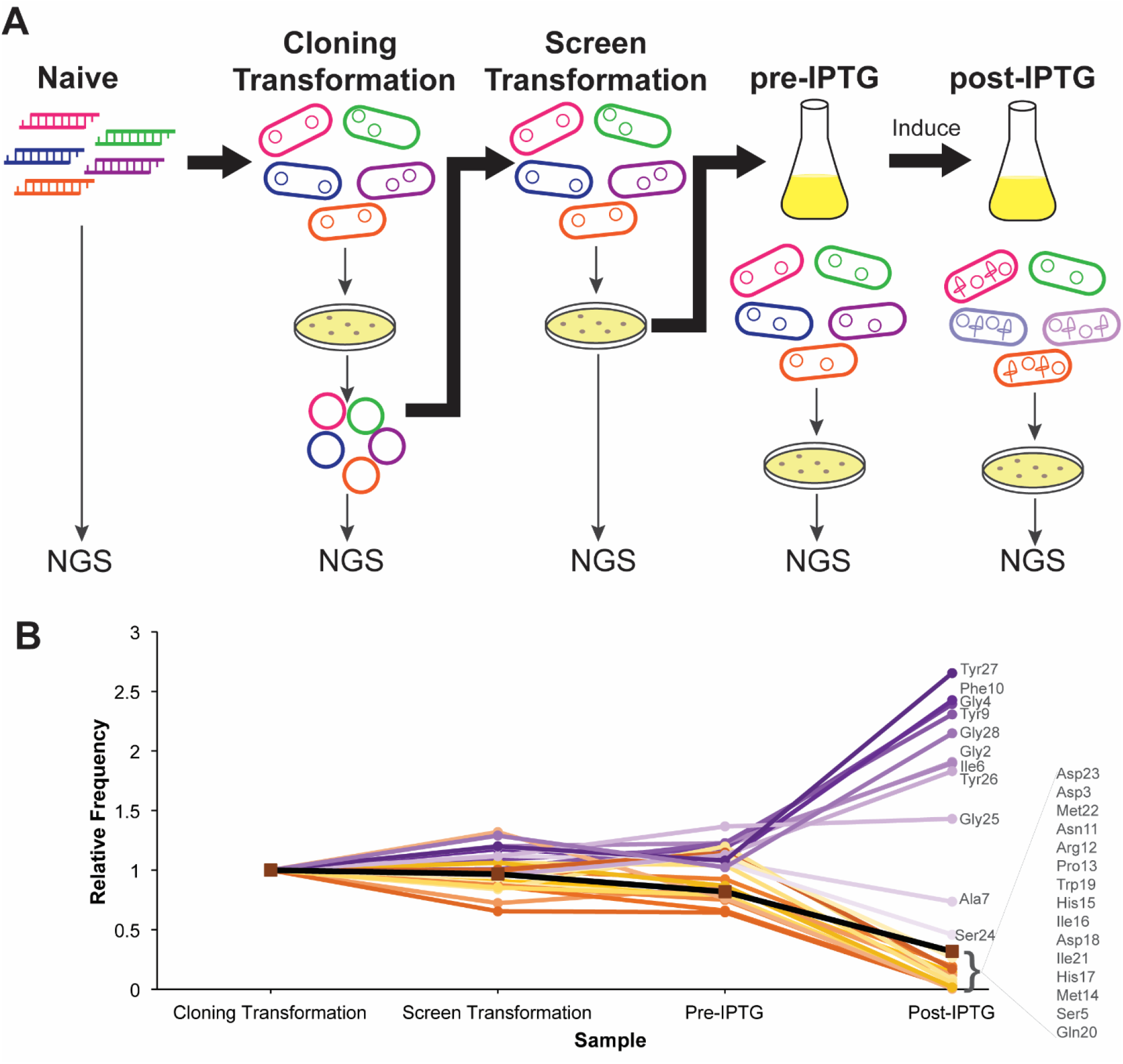
High-throughput screen methodology. **A**: The library was sequenced (NGS, for next gen sequencing) at each of the five stages shown. The naïve library was sequenced as digested PCR products. The cloning transformation was carried out in either XL-1 Blue or DH5*α* while the screen transformation was carried out in MC1061. All samples (except the naïve library) were plated prior to sequencing so that only viable cells were sequenced. **B**: Relative frequencies (Equation S2) averaged over aa substitutions at each residue of ubonodin point variants (excluding mutations to a stop codon) throughout the various steps of the screen. An increase in relative frequency correlates with a loss in RNAP inhibition activity while a drop in relative frequency indicates retention of RNAP inhibition activity. Wild-type (WT) ubonodin decreases in relative frequency and is shown as a reference (squares). Data is from the MiSeq sequencing run and is available as a supplementary file.

First, the naïve library (digested PCR products) was sequenced. These digested PCR products were cloned into the pUboABC backbone and transformed into a *recA*^*-*^ cloning strain. This plasmid library was sequenced again to capture any changes upon transformation of the library into *E. coli*. The change between the naïve library and the cloning transformation library was minimal (Figure S6). This plasmid library was transformed into the screening strain MC1061, and the plasmid from resulting colonies were also sequenced. The library in MC1061 was grown up *en masse* in liquid culture to mid-log phase, and a portion of the culture was plated. These colonies were scraped, then plasmid was isolated and sequenced. Finally, the liquid culture was induced with IPTG after which a portion of the culture was plated and plasmid was isolated and sequenced. It is key to note that all samples from liquid cultures were plated prior to sequencing to ensure that only viable cells were sequenced. Since ubonodin does not lyse cells, directly sampling the liquid culture would cause the plasmids from growth-inhibited and dead cells to be sequenced and the frequencies of ubonodin variants that inhibit RNAP to be artificially inflated. The rationale for sequencing at these five different stages was the possibility that leaky expression of *uboA* would affect the frequency of active mutants prior to IPTG induction. Indeed, we observed that the frequency of some *uboA* mutants dropped as soon as they were transformed into the screening strain MC1061 while the frequency of other mutants climbed (Figure 2B).

### Fitness of Ubonodin Point Variants

The behavior of the point variants (i.e. variants with a single aa substitution) was examined by calculating their enrichment values, the base-2 logarithm of the ratio of the variant’s frequency at a specific step of the screen to the variant’s frequency in the cloning transformation library (Equation S3, Figure S6). The most relevant enrichment value for ubonodin activity is the comparison between the post-IPTG library and the cloning transformation library (Figure 2A). Negative enrichment values indicate that the variant decreased in frequency throughout the screen and likely inhibits RNAP, whereas the opposite is true for positive enrichment values. Many variants inhibit RNAP so strongly that they cannot be detected by NGS after IPTG induction (referred to as dropout variants). After induction with 100 μM IPTG, WT ubonodin has an enrichment value of -4.3. Variants with a stop codon mutation and variants containing an aa substitution at Tyr9 or Tyr27 (positions shown to be crucial for RNAP inhibition in MccJ25) ^26^ have positive enrichment values (1.2 - 1.7), serving as quality control for the screen (Figure 3A). Strikingly, 63% of point variants decreased in frequency, indicating a high tolerance to aa substitution. Hierarchical clustering clearly separates the residues into those with active variants and those with inactive variants (Figure S7A). The positions with active variants are Asn11 through Ser24 in the loop and Asp3, Ser5, and Ala7 in the ring.

**Figure 3.**
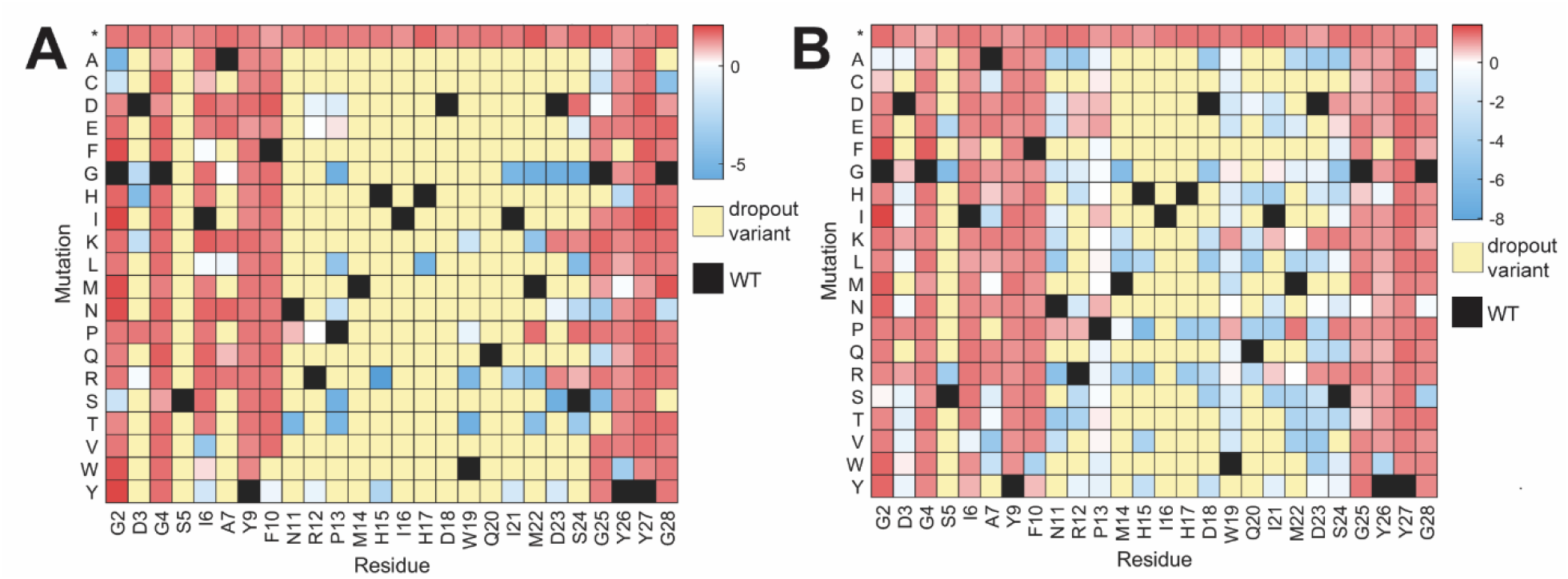
Heatmap of enrichment values for each point variant of ubonodin. Positive enrichment values indicate that the variant does not inhibit RNAP, and negative enrichment values indicate that the variant does inhibit RNAP. Enrichment values after induction of ubonodin production with **A:** 100 μM IPTG, **B:** 10 μM IPTG. Dropout variants were observed in the sequencing runs post induction, likely indicating strong RNAP inhibition. As a reference, the enrichment value for WT ubonodin was -4.3 upon induction with 100 μM IPTG and -4.2 at 10 μM IPTG. Data underlying these heatmaps is from the NovaSeq sequencing run of the single mutant library and is available as a supplementary file.

Induction with 10 μM IPTG serves as a milder condition than induction with 100 μM IPTG (Figures 1C, S4) and led to 56% of point variants decreasing in frequency (Figure 3B). The milder condition shows that not all residues altered within active variants identified earlier are equally potent. The altered residues are separated into three groups: those with the most active variants, those with active variants, and those with inactive variants (Figure S7B). The positions with the most active variants are Met14 through Asp18 and Gln20 in the loop, and Ser5 in the ring. Additionally, substitutions to proline or charged amino acids (lysine, arginine, aspartate, and glutamate) are generally poorly tolerated (Figure S7B).

### Double Mutant Library Construction and Sequencing

We next constructed a focused library of double mutants to exclude variants that likely would not inhibit RNAP. We targeted a subset of ubonodin residues whose point variants tended to retain RNAP inhibitory activity: Asp3, Ser5, and Asn11 through Asp23 (Figure 2B). Sequencing revealed that over 90,000 distinct variants with two aa substitutions were cloned. In addition, all point aa variants were present in the library as were many further variants with three or more aa substitutions were present in the library, as expected from the construction method. To verify the reproducibility of the screen, the enrichment values of point variants in the single mutant screen and point variants in the double mutant screen were compared (Figures 3A, S8). The heatmaps in the two screens are similar with 99% of point variants having the same sign enrichment value in both screens. Next, enrichment values for the double aa variants were computed using the frequency one hour after induction with 100 μM IPTG and plotted on a heatmap in which each axis listed all 520 possible single aa variants (Figure 4). Over 28% of these variants were dropout variants, indicating strong inhibition of RNAP. Hierarchical clustering separates the heatmap into one section with many negative enrichment values and dropout variants (Cluster 1 with 32% of the variants), two sections with many positive enrichment values (Clusters 2 and 3 with 49% of the variants), and one section with many variants absent from the library (Cluster 4 with 19% of the variants). Cluster 1 is largely composed of variants with aa substitutions at two loop residues, though it also contains some variants with aa substitutions at a ring residue (mainly Asp3, Ser5, or Ala7). To analyze the effects of aa substitutions at different residues in the double aa variant library, the spread of enrichment values was examined. Dropout variants were arbitrarily assigned an enrichment value of -20, and the enrichment values were plotted onto a histogram for each residue-specific subset (Figures 5A-B, S9). For example, the Gly4 residue-specific subset contains the enrichment values of all variants with two aa substitutions in which one of the substitutions is at the Gly4 residue. The number of dropout variants differs among the residue-specific subsets and was quantified by calculating the dropout ratio, the base-2 logarithm of the ratio of the number of variants at the mode, (excluding dropout variants) to the number of dropout variants (Figure 5C, Equation S4). Thus, a negative dropout ratio indicates the presence of many distinct dropout variants whereas. Most loop residue-specific subsets have negative dropout ratios, indicating that many of these variants inhibit RNAP strongly.

**Figure 4.**
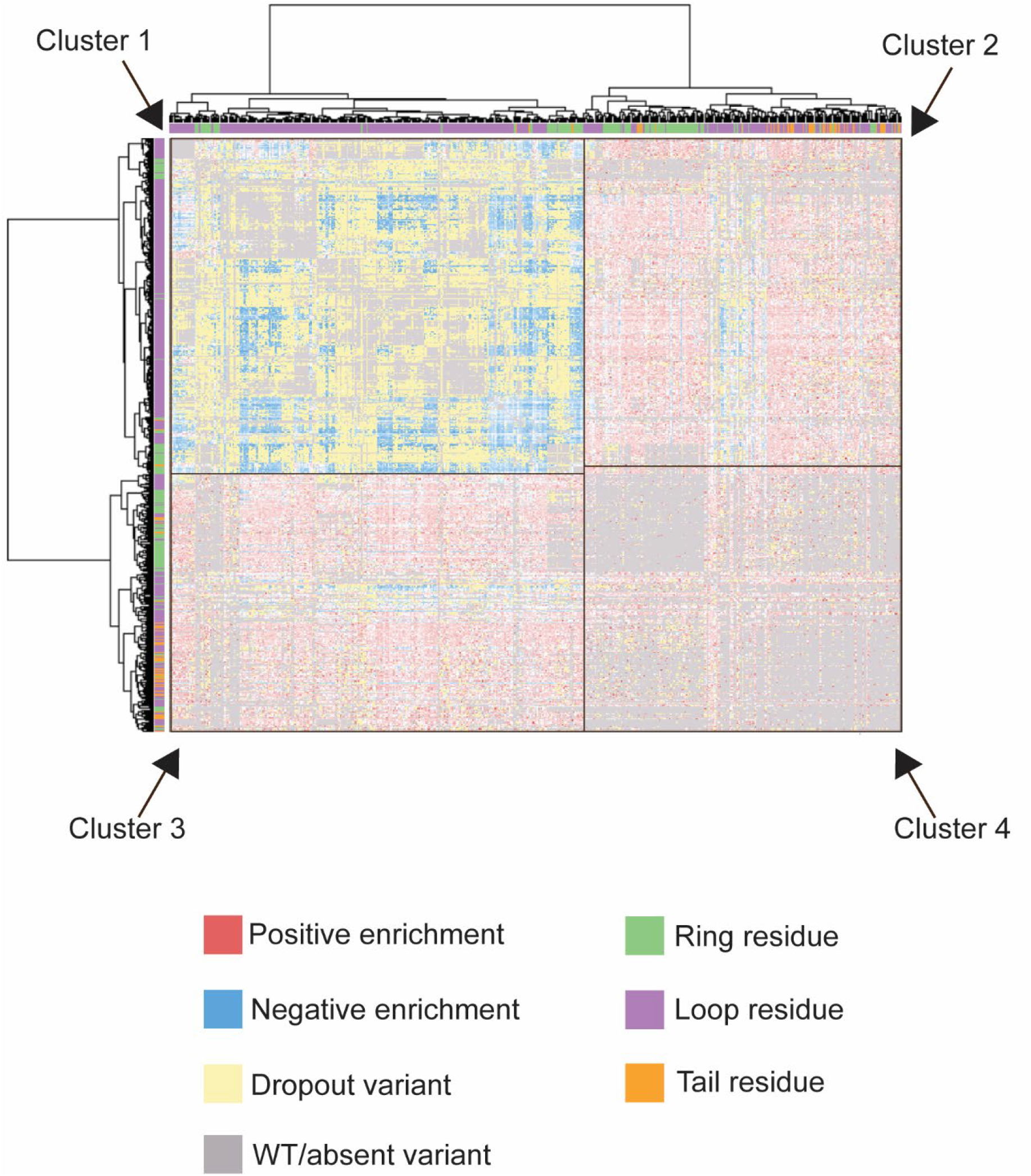
Comprehensive clustered double aa variant heatmap. Each axis lists one possible aa substitution, resulting in 520 entries on each axis. Cluster 1 mainly contains dropout variants and variants with negative enrichment values. Most of the variants in Cluster 1 have two loop residue substitutions. Clusters 2 and 3 mainly contain variants with positive enrichment values. Most of these variants have one substitution in a ring residue or tail residue. Cluster 4 mainly contains absent variants, which either were not targeted for cloning or are variants with substitutions in adjacent residues. The data underlying this heatmap is from the NovaSeq sequencing run of the double mutant library and is available as a supplementary file. An interactive version of the heatmap is also available as a supplementary file.

**Figure 5.**
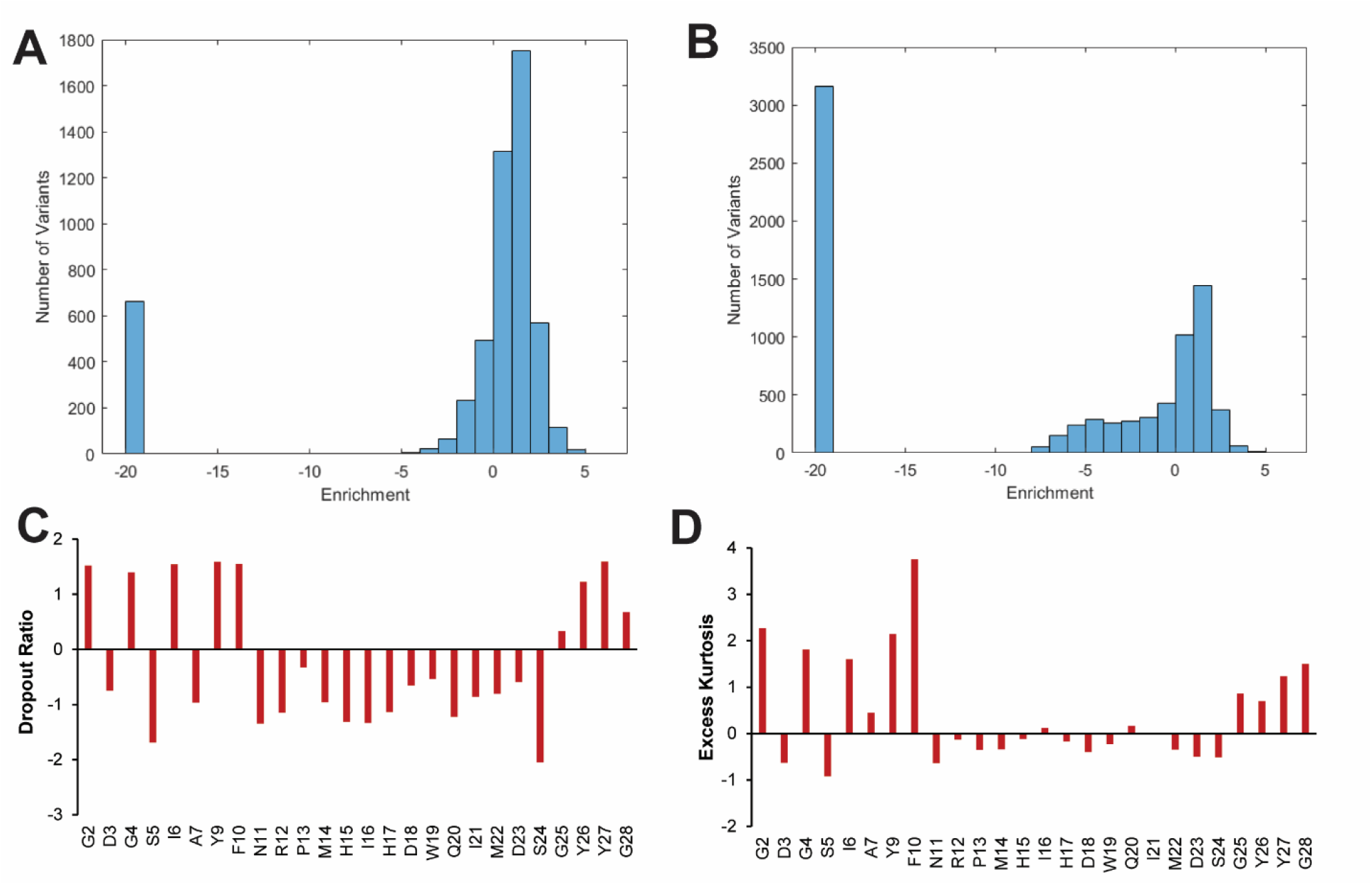
Distributions of enrichment values of residue-specific subsets of the double aa variant library. Histograms of enrichment values for double aa variants in which one aa substitution is at **A**: Gly4, **B:** His17. For double aa variants with a Gly4 substitution, the majority of variants have positive enrichment values corresponding to a loss of RNAP inhibition. The trend is opposite for double aa variants with a substitution at His17; most of these double aa variants retain RNAP inhibition. Histograms for all other positions are in Figure S9. **C**: Dropout ratios of each residue-specific subset. Asp3, Ser5, Ala7, and Asn11 through Ser24 have negative dropout ratios, indicating that substituting these residues leads to a high number of dropout variants corresponding to retention of RNAP inhibition activity. **D**: Excess kurtosis of each residue-specific subset. Gly2, Gly4, Ile6, Tyr9, Phe10, and Gly25 through Gly28 have high excess kurtosis, indicating that a double aa variant with a substitution at one of these residues tends to have enrichment values clustered around the median.

### Rescue Variants

To further examine the behavior of the double aa variants, we calculated excess kurtosis for each residue-specific subset, excluding the dropout variants (Figure 5D, Equation S5). Generally, a positive value describes a narrower distribution than a normal distribution and a negative value describes a wider distribution. A positive excess kurtosis is indicative of a residue-specific subset in which a second aa substitution is unlikely to change the enrichment value. Many ring and tail residue-specific subsets have positive excess kurtosis values (with the median at a positive enrichment value), indicating that for these residues, a second substitution is likely to still result in a variant with a positive enrichment value. However, further examination of the screening data demonstrated the presence of rescue variants which have two aa substitutions in which the second aa substitution changes the enrichment value from positive to negative. In other words, a second beneficial aa substitution rescues the RNAP inhibitory activity of a deleterious first aa substitution. This was observed by manual inspection of non-clustered heatmaps of each residue-specific subset, demonstrating five instances of point variants with rescue variants (Figures 6, S10, Table S1). For example, ubonodin D23R has a positive enrichment value, so most variants in the double aa variant library containing D23R also have positive enrichment values. However, there are 44 D23R double aa variants with negative enrichment values, 14 of which have a substitution at the Ile16 residue. The D23R point variant has an enrichment value of 1.0 but the I16C D23R variant has an enrichment value of -2.9. Strikingly, the residues involved in the rescue variants appear to be physically distal in the structure of ubonodin (Figure 1A). We selected three specific rescue variants and conducted spot dilution assays, experimentally confirming that the second aa substitution does rescue the activity of the first point variant (Figure S11). The presence of these rescue variants could indicate that a deleterious mutation is not necessarily a dead end on the evolutionary trajectory of small peptides like ubonodin.

**Figure 6.**
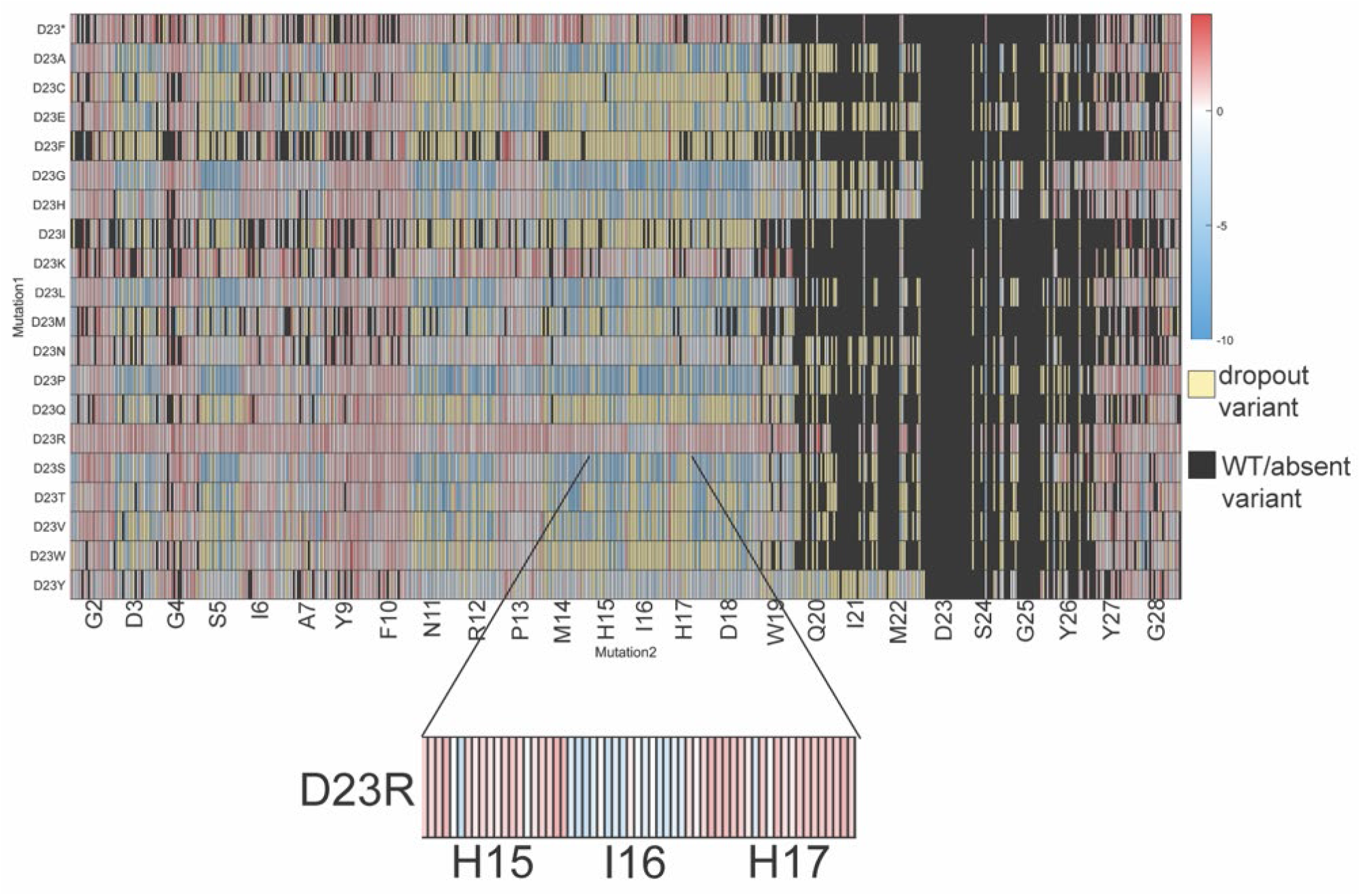
Heatmap of enrichment values for the Asp23 residue-specific subset of the double aa variant library. Most double aa variants containing D23R have positive enrichment values except when the Ile16 residue also has an aa substitution (inset, blue rectangles). In these cases, the second substitution at Ile16 “rescues” the deleterious D23R substitution, restoring some RNAP inhibition activity.

### Deep Learning on the Ubonodin Library: DeepLasso

Since we screened over 90,000 ubonodin variants, we had sufficient data to train a deep learning model, DeepLasso, to predict the enrichment value for ubonodin variants. DeepLasso adopts a classifier-regressor architecture (Figure 7A). With a given input of an ubonodin variant sequence, the classifier first determines whether the variant likely is a dropout variant (with a readout enrichment value of NA). If identified as a non-dropout variant, the regressor is then used to assign an enrichment value to the variant. To represent the ubonodin sequence, DeepLasso was implemented with a sequence encoder to learn the pattern of the ubonodin aa sequence as well as a topology encoder to identify the sequence regions for the ring, loop, and tail of the lasso peptide. The tensors derived from the encoder are concatenated and fed into the classifier for prediction; the resulting tensor from the classifier is then used in the regressor for prediction. Compared with existing deep learning models for prediction of antimicrobial peptides,^37^ the topology encoder we implemented can potentially improve the learning efficiency because the topology of lasso peptides is known to be essential in the inhibition of RNAP.^26, 38-39^

**Figure 7.**
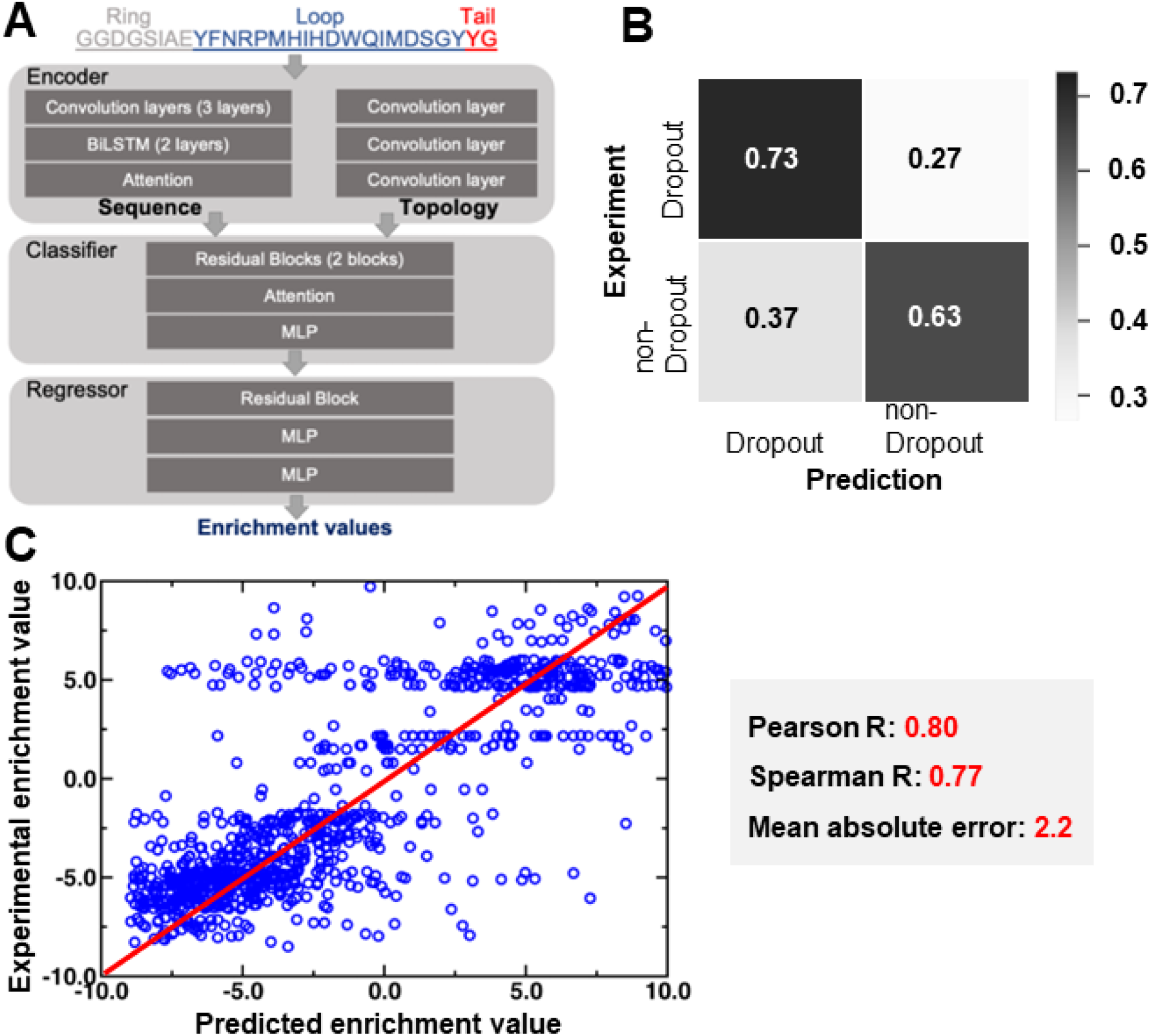
**A**: The architecture of DeepLasso. The encoder consists of a sequence encoder and a topology encoder. The sequence encoder is constructed by three layers of a convolutional neural network, two layers of a bidirectional long-short term memory network (BiLSTM), and one layer of an attention model. The topology encoder is constructed by three layers of a convolutional neural network with each layer used to learn a specific topological part of the lasso peptide (ring, loop, or tail). The classifier involves a sequential layout of two residual blocks, one attention layer, and one layer of multilayer perceptron (MLP). The predictor involves a sequential layout of one residual block and two layers of multilayer perceptron. **B**: Confusion matrix analysis for the classifier of DeepLasso. The matrix shows binary classification of dropout versus non-dropout variants with predicted outcomes on the x-axis and experimental observation on the y-axis. Grayscale is used to represent the magnitude of probability (i.e., high: black; low: white). **C**: Regression analysis for the non-dropout variants with measurable enrichment values. The linear correlation between experimental vs. predicted enrichment values is shown along with Pearson correlation coefficient, Spearman correlation coefficient, and mean absolute error.

The classifier and regressor of DeepLasso were separately trained using five-fold cross-validation with random split (hyperparameters shown in Table S2). The classifier was trained and tested using 61,683 and 6,168 ubonodin sequences, respectively. Both datasets involve two-thirds of dropout variants and one-third of non-dropout variants. The dropout variants were over-represented in the dataset to increase the sensitivity of DeepLasso to identify variants with strong antimicrobial activity. The regressor was trained and tested using 10,330 and 1,033 ubonodin sequences, respectively. All the datasets involve a comprehensive type of mutation, including single, double, triple, quadruple, higher-order, and nonsense mutations (Table S3). This diversity allows DeepLasso to identify potent antimicrobial lasso peptides across large mutational space.

To evaluate the accuracy of DeepLasso, we performed confusion matrix analysis for the classifier (Figure 7B) and linear regression analysis for the regressor (Figure 7C). Based on the test set, DeepLasso achieves a 73% hit rate for the dropout variants and 63% for non-dropout variants (Figure 7B). The higher accuracy for identifying dropout variants is exciting because these variants are the most likely to exhibit strong antimicrobial activity. For non-dropout variants, the predicted enrichment values are correlated to the experimental value with a Pearson correlation R=0.80, a Spearman rank correlation R=0.77, and a mean absolute error of 2.2. The regressor allows us to score the non-dropout variants for their RNAP inhibition activity. For both the classifier and regressor, the accuracy metrics derived from the training set are similar to those from the test sets (Figure S12). This indicates that the predictive models used in DeepLasso are likely not overfitted.

### Antimicrobial Activity of Select Variants against *B. cenocepacia*

We next used our sequencing data to identify the variants with the most potential for antimicrobial activity by calculating each variant’s relative frequency (its frequency relative to the cloning transformation frequency (Equation S2)). We selected seven point variants with monotonic decreasing relative frequencies lower than that of wild-type ubonodin (Figure 8A). However, there were over 12,000 double aa variants that fit the same criteria, so we narrowed our search to double aa variants with greater than 500 reads at the cloning transformation step, restricting the search to ∼1,000 double aa variants. We selected 8 double aa variants such that 6 of them had the same aa substitution as a selected point variant (Figure 8B). These 8 variants included 4 variants that had two substitutions in the loop and 4 variants with one ring and one loop substitution. Though we selected these variants without any guidance from the deep learning study, DeepLasso correctly predicted the RNAP inhibitory activity of the variants in the screen, serving as further validation that the variants were promising hits. There were 4 variants for which DeepLasso did not correctly identify dropout status, but all had very low experimental or predicted enrichment values below -5 (Table S4). Spot dilution assays demonstrated that all selected variants inhibited RNAP (Table 1, Figure S13).

**Table 1.**
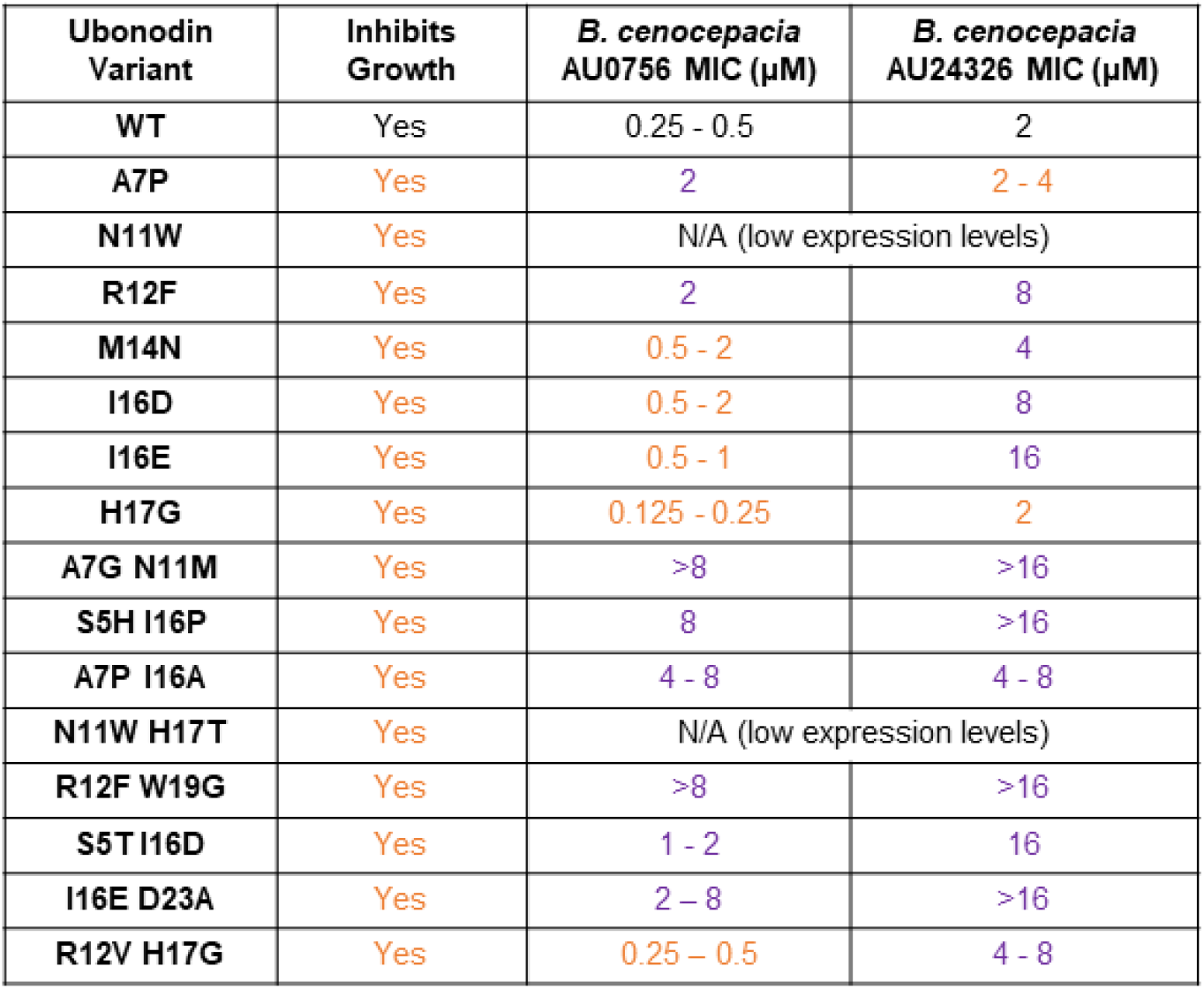
Activity of selected ubonodin variants. Growth inhibition was determined via spot dilution assays (see Figure S13) and MIC was determined via broth microdilution assays. Orange text indicates that the variant has similar or improved activity compared to WT ubonodin and purple text indicates that the variant has decreased activity compared to WT ubonodin.

| Ubonodin Variant | Inhibits Growth | *B. cenocepacia* AU0756 MIC (μM) | *B. cenocepacia* AU24326 MIC (μM) |
| --- | --- | --- | --- |
| WT | Yes | 0.25 - 0.5 | 2 |
| A7P | Yes | 2 | 2 - 4 |
| N11W | Yes | N/A (low expression levels) |  |
| R12F | Yes | 2 | 8 |
| M14N | Yes | 0.5 - 2 | 4 |
| I16D | Yes | 0.5 - 2 | 8 |
| I16E | Yes | 0.5 - 1 | 16 |
| H17G | Yes | 0.125 - 0.25 | 2 |
| A7G N11M | Yes | >8 | >16 |
| S5H I16P | Yes | 8 | >16 |
| A7P I16A | Yes | 4 - 8 | 4 - 8 |
| N11W H17T | Yes | N/A (low expression levels) |  |
| R12F W19G | Yes | >8 | >16 |
| S5T I16D | Yes | 1 - 2 | 16 |
| I16E D23A | Yes | 2 - 8 | >16 |
| R12V H17G | Yes | 0.25 - 0.5 | 4 - 8 |

**Figure 8:**
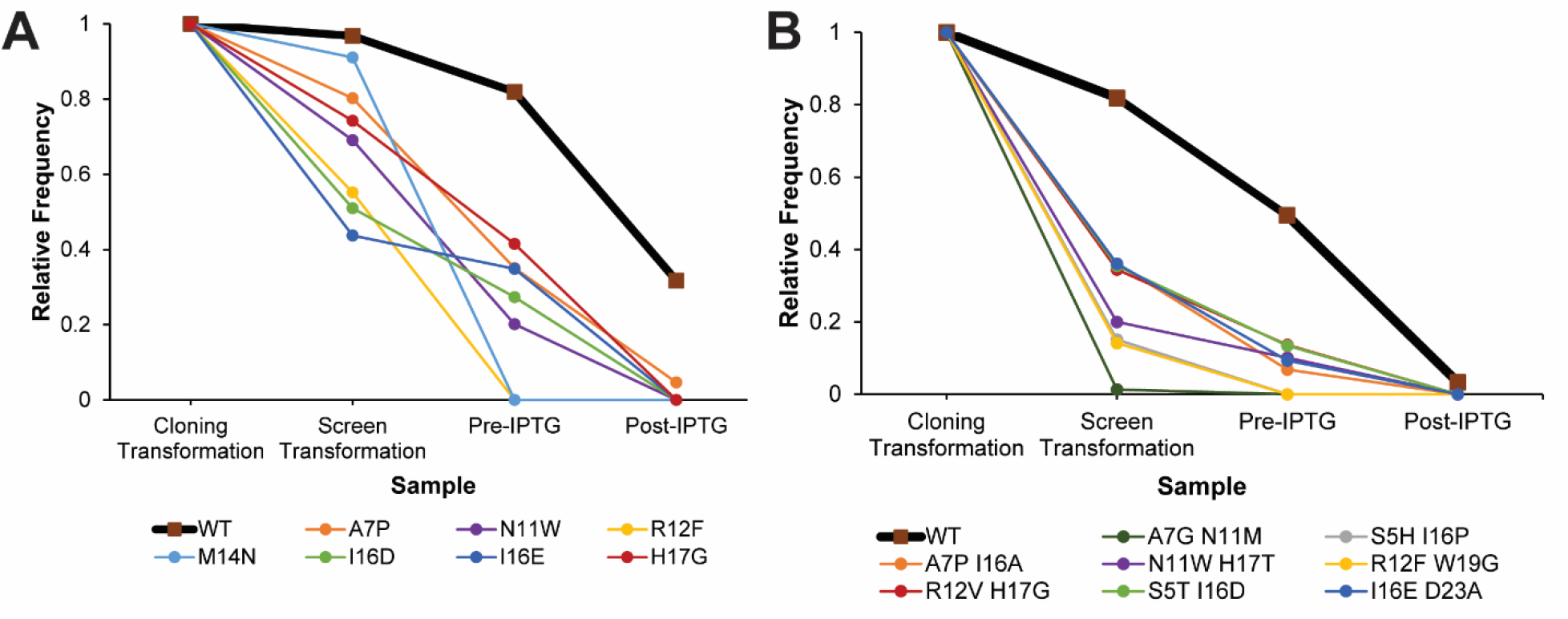
Relative frequencies throughout the screen of selected **A:** ubonodin point variants (data from MiSeq sequencing run), **B**: ubonodin double aa variants (data from NovaSeq sequencing run of the double mutant library). In all cases the relative frequency of the variants decreases more rapidly than that of WT ubonodin.

We proceeded to test the antimicrobial activity of the selected variants against two clinical isolates of the Bcc, *B. cenocepacia* AU0756 and *B. cenocepacia* AU24362, in broth microdilution assays. We have previously reported ubonodin minimal inhibitory concentrations (MIC) of 10 μM against strain AU0756 and 20 μM against strain AU24362 using a spot-on-lawn assay.^36^ We revisited these measurements using broth microdilution assays following the Clinical & Laboratory Standards Institute (CLSI) standard. Ubonodin is more potent in the broth microdilution assay than in the spot-on-lawn assay exhibiting an MIC of 0.25-0.5 μM against strain AU0756 and 2 μM against strain AU24362 (Table 1). Even when accounting for the large mass of ubonodin relative to other antibiotics, the mass basis MIC of ubonodin is competitive with or superior to the MICs of current clinically-deployed antibiotics, including combination therapies (Table S5).^40-42^

Of the 15 ubonodin variants selected, 2 were not tested because of low expression levels. Among the remaining variants, a majority exhibit an MIC within 4-fold of the WT MIC (Table 1). Most notably, the ubonodin H17G variant has an even lower MIC against strain AU0756 than wild-type ubonodin, demonstrating that our screen can identify variants with improved potency. This result is especially remarkable given the already high potency of wild-type ubonodin against *B. cenocepacia* AU0756. Since antimicrobial activity is a product of activity against the target (RNAP) and transport into susceptible cells (through PupB and YddA),^36^ it is not surprising that some of the variants we tested exhibit less potent *B. cenocepacia* inhibition given that transport was not incorporated in our high-throughput screen.

## Discussion

Here we have carried out the most comprehensive structure activity analysis of a lasso peptide to date, with over 90,000 measurements of the fitness of ubonodin variants. From this analysis we found multiple single and double aa variants that are able to inhibit the growth of the Bcc pathogen *B. cenocepacia* at concentrations comparable to that of WT ubonodin. We were even able to identify one variant, H17G ubonodin, that is even more potent than the WT peptide with an MIC of 125 - 250 nM (0.4 - 0.8 μg/mL). The first examination of lasso peptide tolerance to aa substitutions was carried out by Pavlova *et al*. by examining the expression, RNAP inhibition *in vitro*, and antimicrobial activity of a near complete set of 381 point variants of the lasso peptide MccJ25.^3^ Our lab has also previously generated a focused library of triple aa variants of MccJ25, and screened for antimicrobial activity using replica plating with Sanger sequencing of the winners from the screen.^4^ This study revealed structure activity relationships for a further ∼200 MccJ25 variants. Most recently, Hills *et al*. carried out a comprehensive analysis of point variants of the RNAP-inhibiting lasso peptide klebsidin using NGS methodology similar to what we describe here.^5^ This approach has parallels to the Pavlova *et al*. study though it is much more efficient because of the ability to screen and sequence the library *en masse*. The picture that emerges from these studies is that the ability of a lasso peptide to absorb aa substitutions while still retaining WT-like activity depends on the peptide being mutagenized. Whereas multiple variants of MccJ25 and ubonodin could be obtained with near or better activity than the WT peptide, nearly all aa substitutions to klebsidin were deleterious to its activity. This difference may be due to the fact that klebsidin is inherently less potent than MccJ25 and ubonodin.

Our analysis of ubonodin double aa variants also allowed for deeper insights than was possible with the single aa or focused library studies described above. First, our data suggest that ubonodin variants with RNAP inhibition activity comparable to that of the wild-type peptide are fairly common within sequence space (Figure 4). The dataset also revealed rescue variants in which a 2^nd^ aa substitution is able to rescue an otherwise deleterious primary substitution, converting a non-RNAP inhibiting ubonodin variant back to one that can inhibit RNAP. These results give further insights into the sequence/function space of ubonodin. The large dataset about the fitness of ubonodin variants also allowed for the development of DeepLasso, a deep learning approach to predict the RNAP inhibition capability of an ubonodin variant. While other deep learning packages have been developed for membrane-active antimicrobial peptides,^37^ this work represents the first time that deep learning approaches have been applied to lasso peptides. Beyond predicting the RNAP inhibitory activity of double aa variants, DeepLasso will have utility in scoring triple, quadruple, and further aa variants for which the construction and experimental validation of the entire library is infeasible. We also plan to test whether the lessons learned from deep learning on ubonodin can predict activity of other RNAP-inhibiting lasso peptides.

One limitation of our approach was revealed when testing highly-ranked variants from our screen against *B. cenocepacia* directly. Some of these variants that appear to inhibit RNAP well when produced inside the cell are only mediocre in the broth microdilution assay. This suggests that some substitutions that may be beneficial (or neutral) for RNAP inhibition are deleterious for the transporters that internalize ubonodin in susceptible cells. In the future, we plan to utilize this dataset and DeepLasso to construct focused libraries that can be screened directly against Bcc members, perhaps by using a platform akin to nanoFleming.^6^ Such a screen would provide us with information on variants that can be transported into susceptible Bcc, which could be used to further train DeepLasso to predict potent variants. This effort will ultimately lead to ubonodin variants that can serve as leads for new antibiotics to treat Bcc infections.

## Acknowledgements

We thank Wei Wang at the Lewis-Sigler Institute for Integrative Genomics at Princeton University for guidance with preparing and processing our NGS samples. We also thank Dr. John J. LiPuma at University of Michigan Pediatrics Infectious Disease for sharing the *B. cenocepacia* AU0756 and AU24326 strains with us. This work was supported by the National Institutes of Health Grant GM107036 and a grant from Princeton University School of Engineering and Applied Sciences (Focused Research Team on Precision Antibiotics). Z.Y. and R. X. were supported by Vanderbilt University and a scholarship from the Vanderbilt Institute of Chemical Biology. This work was carried out in part using computational resources from the Extreme Science and Engineering Discovery Environment (XSEDE), which is supported by National Science Foundation grant number TG-BIO200057.

## Methods Summary

Full detailed methods are in the Supporting Information

### Cloning

Plasmids were constructed using overlap PCR, restriction digestion, and ligation followed by transformation into *E. coli* XL1-Blue. Plasmid sequences were verified using Sanger sequencing. Additional information regarding plasmid construction can be found in the Supporting Information.

### Spot Dilution Assays

Cultures of *E. coli* MC1061 transformed with the appropriate plasmids were grown. Aliquots of the cultures were sampled prior to and after induction with 100 μM IPTG, and ten-fold serial dilutions were plated.

### Colony-Forming Units of *E. coli* pUboABC MC1061

*E. coli* MC1061 pUboABC and *E. coli* MC1061 pUboBC cultures were grown. Aliquots of the cultures were sampled prior to and after induction with 10 μM IPTG and 100 μM IPTG, and ten-fold serial dilutions were plated. Colonies were counted 16 hours after incubation.

### Co-Culture of *E. coli* MC1061 pUboABC and *E. coli* MC1061 pUboBC

A culture containing approximately 90% *E. coli* MC1061 pUboABC and 10% *E. coli* MC1061 pUboBC was grown. Aliquots of the culture were sampled prior to and after induction with 10 μM IPTG and 100 μM mM IPTG, and ten-fold serial dilutions were plated. Sixteen hours after incubation, colonies were resuspended in media and plasmids were miniprepped and PCR amplified with primers that amplified regions of different sizes in pUboABC and pUboBC. Amplicons were visualized on an agarose gel.

### Library Construction

Site-saturation mutagenesis libraries were constructed with overlap PCR using mutagenic NNK primers, restriction digestion, and ligation. pWC99 was used as the PCR template for constructing the single mutant library, and the single mutant library was used as the PCR template for constructing the double mutant library. Transformations yielded at least 10-fold coverage of all possible nucleotide mutants.

### Screen Methodology

The single mutant and double mutant libraries were transformed into *E. coli* MC1061. Colonies were resuspended in media and sub-cultured into a larger culture. Aliquots of the cultures were sampled prior to and after inducing with 10 μM IPTG and 100 μM IPTG and plated. After 16 hours of incubation, the colonies were resuspended in media and plasmids were miniprepped.

### NGS

Samples were PCR amplified with barcoded primers and combined such that double mutant library samples were over-represented compared to single mutant library samples to provide sufficient coverage of the larger double mutant library. Samples were sequenced using an Illumina MiSeq Micro 300nt and an Illumina NovaSeq 6000 Sequencing System. Downstream analysis of the data was conducted with Python, MATLAB, and R.

### Expression and Purification of Ubonodin Variants

All ubonodin variants were expressed in *E. coli* BL21 cells and induced with 1 mM IPTG. Cultures were centrifuged and supernatants were applied to a C8 column prior to continuing purification with HPLC. Purity was verified with an LC-MS.

### Antimicrobial Assays

Broth microdilution assays of the ubonodin variants were conducted using *B. cenocepacia* AU0756 and *B. cenocepacia* AU24362 with guidelines provided by the Clinical & Laboratory Standards Institute (CLSI).

## Synopsis

We screened a large library of variants of the lasso peptide ubonodin for their antimicrobial activity, allowing us to identify ubonodin variants with potent activity.

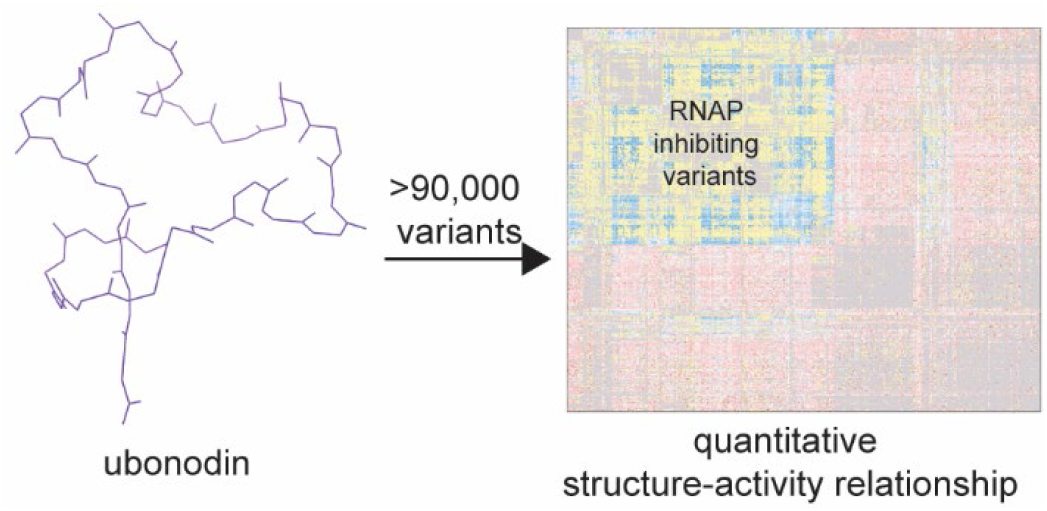

