## Supporting Info for "A High-Throughput Screen Reveals the Structure-Activity Relationship of the Antimicrobial Lasso Peptide Ubonodin"

**Supporting Information**

Alina Thokkadam[1], Truc Do[1], Xinchun Ran[2], Mark P. Brynildsen[1,3], Zhongyue J. Yang[2,4-6], A. James Link[1,3,7,*]

[1]Department of Chemical and Biological Engineering, Princeton University, Princeton, NJ 08544, United States

[2]Department of Chemistry, Vanderbilt University, Nashville, TN 37235, United States

[3]Department of Molecular Biology, Princeton University, Princeton, NJ 08544, United States

[4]Department of Chemical and Biomolecular Engineering, Vanderbilt University, Nashville, TN 37235, United States

[5]Data Science Institute, Vanderbilt University, Nashville, TN 37235, United States

[6]Vanderbilt Institute of Chemical Biology, Vanderbilt University, Nashville, TN 37235, United States

[7]Department of Chemistry, Princeton University, Princeton, NJ 08544, United States

**Table of Contents**

Methods…………………………………………………………..…...……………...…….. Pages 3-9

Supporting Figures……………...………………………………………………………...Pages 10-29

Supporting Tables…………………………………………………………………………..Pages 30-43

Supporting References…………………………………………………………………..Page 44

Methods

### Safety

No unexpected or unusually high safety hazards were encountered.

### Cloning

All cloning was conducted by PCR amplification (using Q5 polymerase), restriction digestion, ligation, (using T4 DNA ligase), and transformation into chemically competent *E. coli* XL1-Blue. All cloning with a pUboABC backbone was done by plating transformation mixtures onto Lysogeny Broth (LB) plates supplemented with 100 µg/mL ampicillin and 0.2% w/v glucose, and colonies were inoculated into LB media supplemented with 100 µg/mL ampicillin and 0.2% w/v glucose to suppress any leaky expression. Cloning for all other plasmid constructs was done by plating transformation mixtures onto Lysogeny Broth (LB) plates supplemented with 100 µg/mL ampicillin, and colonies were inoculated into LB media supplemented with 100 µg/mL ampicillin. Plasmids were extracted using Qiagen QIAprep Spin Miniprep Kit and sequence verified using Sanger sequencing. All primer sequences are listed in Table S6, and all plasmid constructs are listed in Table S7.

pUboABC was cloned by amplifying pWC99 with pQE-80 *Nhe*I For and pAT8 *Nco*I Rev. The amplicon was ligated into pWC99 digested with *Nhe*I and *Nco*I.

pUboBC was cloned by digesting pUboABC with *Nhe*I and *Nco*I and ligating that fragment into pQE-80 digested with *Nhe*I and *Nco*I.

pUboABC S5H I16P and pUboABCD S5H I16P were cloned with overlap PCR using pWC99 as the template. One fragment was amplified with pQE-80 *Eco*RI For and S5H I16P Rev v2 and the second fragment was amplified with S5H I16P For v2 and pQE-80 *Hin*dIII Rev. The fragments were joined with a second round of PCR and was amplified with pQE-80 *Eco*RI For and pQE-80 *Hin*dIII Rev. pUboABC S5H I16P was cloned by ligating the amplicon into pUboBC digested with *Eco*RI and *Hin*dIII. pUboABCD S5H I16P was cloned by ligating the insert into pWC99 digested with *Eco*RI and *Hin*dIII.

All remaining plasmid constructs were cloned with overlap PCR. One fragment was amplified with *Xho*I For Lib and a mutagenic reverse primer and the second fragment was amplified with *Nhe*I Rev Lib and a mutagenic forward primer. The templates and mutagenic primers for each plasmid construct are listed in Table S7. The fragments were joined with a second round of PCR and amplified with *Xho*I For Lib and *Nhe*I Rev Lib. Amplicons were ligated into the appropriate plasmid backbone digested with *Xho*I and *Nhe*I-HF. The plasmid backbones used for each plasmid construct are listed in Table S7.

### Spot Dilution Assay

The appropriate plasmid was transformed into electrocompetent *E. coli* MC1061, plated onto LB plates supplemented with 100 µg/mL ampicillin and 0.2% w/v glucose, and incubated at 30 °C for 16 hours. Colonies were inoculated into LB media supplemented with 100 µg/mL ampicillin and 0.2% w/v glucose and grown at 37 °C for 16 hours. These cultures were sub-cultured to an $OD_{600}$ of 0.02 in 50 mL of LB media supplemented with 100 µg/mL ampicillin in a 250 mL flask. Once the $OD_{600}$ reached 0.3, a 1 mL aliquot was sampled immediately prior to inducing the cultures with 100 µM isopropyl β-D-1-thiogalactopyranoside (IPTG). The aliquot was washed with Phosphate-Buffered Saline (PBS) two times and ten-fold serial dilutions were spotted onto LB plates supplemented with 100 µg/mL ampicillin and 0.2% w/v glucose. For each following timepoint, a 1 mL sample of each culture was withdrawn, washed with PBS two times, and ten-fold serial dilutions were spotted onto LB plates supplemented with 100 μg/mL ampicillin and 0.2% w/v glucose. Plates were incubated for 16 hours at 37 °C before imaging.

**Colony-Forming Units of *E. coli* MC1061 pUboABC and *E. coli* MC1061 pUboBC**

pUboABC and pUboBC were transformed into electrocompetent *E. coli* MC1061, plated onto LB plates supplemented with 100 μg/mL ampicillin and 0.2% w/v glucose, and incubated at 37 °C for 16 hours. Single colonies were inoculated into LB media supplemented with 100 μg/mL ampicillin and 0.2% w/v glucose and shaken at 37 °C for 16 hours. These cultures were then sub-cultured to an $OD_{600}$ of 0.02 into 50 mL of LB media supplemented with 100 μg/mL ampicillin in a 250 mL flask. Ten-fold serial dilutions of the culture were plated onto LB plates supplemented with 100 μg/mL ampicillin and 0.2% w/v glucose. The cultures were then grown at 37 °C, and once the $OD_{600}$ reached 0.3, a 1 mL sample of the culture was withdrawn immediately before the cultures were induced with 10 μM or 100 μM IPTG and returned to grow longer. The 1 mL sample was washed with PBS two times and ten-fold serial dilutions were plated onto LB plates supplemented with 100 μg/mL ampicillin and 0.2% w/v glucose. For each following timepoint, a 1 mL sample of each culture was withdrawn, washed with PBS two times, and ten-fold serial dilutions were plated onto LB plates supplemented with 100 μg/mL ampicillin and 0.2% w/v glucose. All plates were incubated at 37 °C for 16 hours, after which the number of colonies was counted. Three biological replicates of this assay were done.

**Co-Culture of *E. coli* MC1061 pUboABC and *E. coli* MC1061 pUboBC**

pUboABC (0.9 ng) and pUboBC (0.1 ng) were transformed into electrocompetent *E. coli* MC1061 with a one-hour outgrowth at 37 °C in LB media supplemented with 0.2% w/v glucose. A small quantity of DNA was used to reduce the probability of two plasmids being transformed into the same cell. The transformation mixture was plated onto an LB plate supplemented with 100 μg/mL ampicillin and 0.2% w/v glucose and incubated at 37 °C for 16 hours, yielding at least 1,000 colonies. Colonies were resuspended in LB media supplemented with 100 μg/mL ampicillin and sub-cultured to an $OD_{600}$ of 0.02 into 50 mL of LB media supplemented with 100 μg/mL ampicillin in a 250 mL flask. Plasmids were extracted from the remaining resuspended colony mixture using a Qiagen QIAprep Spin Miniprep Kit and labeled as the MC1061 transformation sample.

The culture was grown until $OD_{600}$ reached 0.3 at which point 1 mL samples were aliquoted and washed with PBS two times prior to plating dilutions onto LB plates supplemented with 100 μg/mL ampicillin and 0.2% w/v glucose. The culture was then split into two flasks of 25 mL each; one flask was induced with 10 μM IPTG and another flask was induced with 100 μM IPTG. Subsequent timepoints were taken by aliquoting 1 mL samples from the cultures, washing with PBS two times, and plating dilutions onto LB plates supplemented with 100 μg/mL ampicillin and 0.2% w/v glucose. Plates were incubated at 30 °C for 16 hours; each sample had at least 10,000 colonies. Colonies were resuspended in LB media supplemented with 100 μg/mL ampicillin and 0.2% w/v glucose, and plasmids were extracted using a Qiagen QIAprep Spin Miniprep Kit. Each plasmid sample, as well as a sample of 90% pUboABC and 10% pUboBC, was PCR amplified. Each PCR reaction contained 100 ng of template DNA and was amplified with *Xho*I For and *Nhe*I Rev primers for 10 cycles. Reactions were then run on a 1.5% agarose gel. Two biological replicates of this assay were done.

**Library Construction: Single Mutants**

Single-site saturation mutagenesis (SSM) libraries were created with overlap PCRs using Q5 polymerase for each of the 26 mutated positions in the ubonodin core peptide (all positions excluding the Gly1 and Glu8 residues). The PCR template was pWC99 treated with Exonuclease

V to reduce genomic DNA contamination. One fragment was amplified with *Xho*I For Lib and a mutagenic reverse primer and the second fragment was amplified with *Nhe*I Rev Lib and a mutagenic forward primer. The mutagenic primers for each residue-specific library are listed in Table S8. A low annealing temperature of 55 °C was used to promote binding of mutagenic primers. The fragments were joined with a second round of PCR and amplified with *Xho*I For Lib and *Nhe*I Rev Lib. A residue-specific library hereafter refers to the portion of the library that was cloned with one set of mutagenic primers. For example, the G2 residue-specific library refers to the portion of the library that was cloned using the G2 Lib For and G2 Lib Rev primers.

Amplicons were digested with *Nhe*I-HF and *Xho*I using restriction digestion and ligated into pUboABC that had been digested with *Nhe*I-HF and *Xho*I and treated with Antarctic Phosphatase. Digested inserts were stored at -20 °C for next-generation sequencing (NGS). Ligation mixtures were directly transformed into chemically competent *E. coli* XL1-Blue, plated onto LB plates supplemented with 100 µg/mL ampicillin and 0.2% w/v glucose, and incubated at 37 °C for 16 hours, leading to at least 320 colonies per residue mutated (yielding 10-fold coverage of all nucleotide mutants). Colonies were subsequently resuspended in 5 mL of LB media supplemented with 100 µg/mL ampicillin and plasmids were extracted using a Qiagen QIAprep Spin Miniprep Kit. All residue-specific libraries were cloned separately, then pooled at equimolar levels and sequenced with an Illumina MiSeq Micro 300nt. Data on the pooled samples indicated that a few G2 variants were missing (pUboABC G2F, G2I, G2M, G2N, G2W, and G2Y), so plasmids encoding these variants were individually cloned, then added to the library such that all residue-specific libraries were still present at roughly equimolar levels. This sample is hereafter referred to as the single mutant library.

**Library Construction: Double Mutants**

Single-site saturation mutagenesis (SSM) libraries were created with overlap PCRs using Q5 polymerase for each of the ubonodin core peptide positions to be mutated (at residues Asp3, Ser5, Asn11, Arg12, Pro13, Met14, His15, Ile16, His17, Asp18, Trp19, Gln20, Ile21, Met22, and Asp23). The PCR template was the single mutant library treated with Exonuclease V to reduce genomic DNA contamination. One fragment was amplified with *Xho*I For Lib and a mutagenic reverse primer and the second fragment was amplified with *Nhe*I Rev Lib and a mutagenic forward primer. The mutagenic primers for each residue-specific library are listed in Table S8. A low annealing temperature of 55 °C was used to promote binding of mutagenic primers. The fragments were joined with a second round of PCR and amplified with *Xho*I For Lib and *Nhe*I Rev Lib. Amplicons were digested with *Nhe*I-HF and *Xho*I using restriction digestion and ligated into pUboABC that had been digested with *Nhe*I-HF and *Xho*I and treated with Antarctic Phosphatase. Digested inserts were stored at -20 °C for NGS. Ligation mixtures were then desalted using a 0.025 µM nitrocellulose membrane in water. Desalted ligation mixtures were transformed into freshly prepared electrocompetent *E. coli* DH5α with a one-hour outgrowth in Super Optimal broth with Catabolite repression (SOC) at 37 °C. Transformation mixtures were plated onto LB plates supplemented with 100 µg/mL ampicillin and 0.2% w/v glucose, and incubated at 30 °C for 16 hours, yielding at least 120,000 colonies per residue mutated (yielding 10-fold coverage of all nucleotide mutants). Colonies were subsequently resuspended in 15 mL of LB media supplemented with 100 µg/mL ampicillin, and plasmids were extracted using a Qiagen QIAprep Spin Miniprep Kit. Residue-specific libraries were then combined in equimolar amounts, constituting the double mutant library.

**Screen Methodology: MiSeq Sequencing**

One ng of the single mutant library was transformed into freshly prepared electrocompetent *E. coli* MC1061 cells with a one-hour outgrowth in LB media supplemented with 0.2% w/v glucose at 37 °C. A small quantity of DNA was used to reduce the probability of two plasmids being transformed into the same cell. Transformation mixtures were plated onto LB plates supplemented with 100 µg/mL ampicillin and 0.2% w/v glucose and incubated at 30 °C for 16 hours to obtain ~3.7 x $10^5$ colonies (at least 8 x $10^3$ colonies were needed to obtain at least 10-fold coverage of all nucleotide mutants). Colonies were resuspended in LB media supplemented with 100 µg/mL ampicillin. Each library was sub-cultured to an $OD_{600}$ of 0.02 into a 250 mL flask containing 50 mL of LB media supplemented with 100 µg/mL ampicillin. Plasmids were extracted from the remaining resuspended colony mixture using a Qiagen QIAprep Spin Miniprep Kit and labeled as the screen transformation sample.

The 50 mL cultures were grown at 37 °C until the $OD_{600}$ reached 0.3. A 1 mL aliquot was sampled from the culture, washed with PBS two times, and plated onto LB plates supplemented with 100 µg/mL ampicillin and 0.2% w/v glucose. Immediately after aliquoting, the culture was induced with 100 µM IPTG and returned to the shaker. One hour after induction, a 1 mL sample was aliquoted from the culture, washed with PBS two times, and dilutions were plated onto LB plates supplemented with 100 µg/mL ampicillin and 0.2% w/v glucose. The plates were incubated at 30 °C for 16 hours, yielding at least 4.9 x $10^4$ colonies for all timepoints (at least 8 x $10^3$ colonies were needed to obtain at least 10-fold coverage of all nucleotide mutants). The colonies from each plate were resuspended in 15 mL of LB media supplemented with 100 µg/mL ampicillin, and plasmids were extracted using a Qiagen QIAprep Spin Miniprep Kit.

**Screen Methodology: NovaSeq Sequencing**

One ng each of the single mutant library and double mutant library were transformed into freshly prepared electrocompetent *E. coli* MC1061 cells with a one-hour outgrowth in LB media supplemented with 0.2% w/v glucose at 37 °C. A small quantity of DNA was used to reduce the probability of two plasmids being transformed into the same cell. Transformation mixtures were plated onto LB plates supplemented with 100 µg/mL ampicillin and 0.2% w/v glucose and incubated at 30 °C for 16 hours to obtain ~1.4 x $10^6$ colonies for the single mutant library and ~2.1 x $10^6$ colonies for the double mutant library (at least 8 x $10^3$ colonies were needed for the single mutant library and at least 2 x $10^6$ colonies were needed for the double mutant library to obtain at least 10-fold coverage of all nucleotide mutants). Colonies were resuspended in LB media supplemented with 100 µg/mL ampicillin. Each library was sub-cultured to an $OD_{600}$ of 0.02 into a 250 mL flask containing 50 mL of LB media supplemented with 100 µg/mL ampicillin. Plasmids were extracted from the remaining resuspended colony mixture using a Qiagen QIAprep Spin Miniprep Kit and labeled as the screen transformation sample.

The 50 mL cultures were grown at 37 °C until the $OD_{600}$ reached 0.3. One mL aliquots were sampled from each culture, washed with PBS two times, and plated onto LB plates supplemented with 100 µg/mL ampicillin and 0.2% w/v glucose. Each culture was then split into two 250 mL flasks with 25 mL of culture in each; one flask was induced with 10 µM IPTG and another flask was induced with 100 µM IPTG. The following timepoints were taken: 1 hour after induction for both induction conditions, and 2 hours and 3 hours after induction for the 10 µM IPTG condition. The timepoints were taken by aliquoting 1 mL samples from the cultures, washing with PBS two times, and plating dilutions onto LB plates supplemented with 100 µg/mL ampicillin and 0.2% w/v glucose. The plates were incubated at 30 °C for 16 hours, yielding at least 4.2 x $10^4$ colonies for all timepoints for the single mutant library and at least 7.8 x $10^6$ colonies for all time points for the double mutant library (at least 8 x $10^3$ colonies were needed for the single mutant library and at least 2 x $10^6$ colonies were needed for the double mutant library to obtain at least 10-fold coverage of all nucleotide mutants). The colonies from each plate were resuspended in 15 mL of LB media supplemented with 100 µg/mL ampicillin, and plasmids were extracted using a Qiagen QIAprep Spin Miniprep Kit.

### Next-Generation Sequencing: Library Preparation

Each sample was PCR amplified with Q5 polymerase and 10 amplification cycles. Primers amplified between the Gly2 and Gly28 DNA region of the plasmids. Both forward and reverse primers had an adaptor sequence. Reverse primers (all primers in Table S6 that begin with "P7") also had a barcode specific to each sample. The forward primer, P5-AT1F, was used for all samples but the reverse primers were different for each sample. The primer sequences are listed in Table S6. The DNA was purified using a Zymo DNA Clean & Concentrator. The single mutant library was initially sequenced using an Illumina MiSeq Micro 300nt. When sequencing on the Illumina NovaSeq 6000 Sequencing System, the samples from the screen were combined such that each single mutant library sample represented 1% of the total DNA and each double mutant library sample represented 11.5% of the total DNA to achieve sufficient coverage of the larger double mutant library.

### Next-Generation Sequencing: Processing Raw Data

The sequencing results were demultiplexed using Barcode Splitter 0.18.4.0 in Galaxy. The demultiplexed DNA sequencing reads in FASTA format were further processed on the Princeton Della server using the custom Python code NGS-code-annotated.py that can be found at the Link lab Github page: https://github.com/ajlinklab/Ubolib . This code iterated through each unique DNA sequence that was read and directionally compared it to the wild-type ubonodin DNA sequence (5'-GGCGATGGCAGCATTGCGGAATACTTTAACCGTCCGATGCATATTCATGATTGGCAGATTATGGATAGCGGCTATTATGGC-3') and the wild-type ubonodin amino acid sequence (N-GDGSIAEYFNRPMHIHDWQIMDSGYYG-C) to identify any mutations and their locations. Due to trimming of sequencing reads, note that the DNA sequence starts at the second codon and the amino acid sequence starts at the second amino acid. For comparison of amino acid sequences, the original DNA sequence that was read was first translated using the standard codon table within the Biopython package. Note that if multiple mutations were found in a single amino acid sequence, the mutations and their counts were recorded as a single entity (e.g., the S5T E8D double mutant appears 140 times in total across the reads found in the current FASTA file). By contrast, comparison of the DNA sequence was conducted on a codon-by-codon basis and multiple codon mutations that were found in a single DNA sequence were recorded as separate entities (e.g., the GGC10GTC ATT16TTT double mutant is recorded as GGC7GTC and ATT13TTT each appearing 89 times in total across the reads found in the current FASTA file). Identified mutations and their counts were saved as tab-delimited output text files. The data is deposited in NCBI as BioProject number PRJNA894114.

### Next-Generation Sequencing: Additional Data Analysis

For the single mutant library samples sequenced with the NovaSeq, all amino acid variants with >500 reads were kept, and for the double mutant library samples sequenced with the NovaSeq, all amino acid variants with 10 reads or higher were kept. For the single mutant library sequenced samples with the MiSeq, all amino acid variants with 40 reads or higher were kept, except for the post-IPTG sample in which all amino acid variants with 10 reads or higher were kept. The frequency of each variant in each sample was calculated using Equation S1.

Equation S1: $frequency = \frac{Number\ of\ reads\ of\ variant}{Total\ number\ of\ reads\ in\ sample} \times 100\%$

Relative frequencies were calculated to compare the change in frequency of a variant throughout the screen using Equation S2. Line graphs were constructed using relative frequency values and plotted using Excel.

Equation S2: $Relative\ frequency = \frac{Frequency\ of\ variant\ in\ sample}{Frequency\ of\ variant\ at\ cloning\ transformation}$

The enrichment was calculated to compare the change in frequency of a variant from the cloning transformation to another sample using Equation S3. Variants that increased in frequency had positive enrichment values and variants that decreased in frequency had negative enrichment values.

Equation S3: $enrichment = \log_2 \frac{frequency\ of\ variant\ [specific\ sample]}{frequency\ of\ variant\ [cloning\ transformation]}$

Histograms were constructed using MATLAB. The dropout ratio was calculated using Equation S4. Excess kurtosis was calculated in MATLAB using Equation S5 where $\mu$ is the mean of $x$, $\sigma$ is the standard deviation of $x$, and $E(t)$ represents the expected value of the quantity $t$.

Equation S4: $dropout\ ratio = \log_2 \frac{Number\ of\ variants\ at\ mode\ (excluding\ dropout\ variants)}{Number\ of\ dropout\ variants}$

Equation S5: $excess\ kurtosis = \frac{E(x-\mu)^4}{\sigma^4} - 3$

Frequency of residue-specific variants were determined for the single mutant library samples using Equation S6:

Equation S6: $frequency\ of\ residue\ variants = \frac{\sum Reads\ for\ all\ variants\ of\ a\ residue}{Total\ number\ of\ reads\ in\ sample} \times 100\%$

Non-clustered heatmaps were constructed using MATLAB. In order for the dropout variants to affect the clustering in the clustered heatmaps, the dropout variants in the clustered single mutant heatmaps were arbitrarily assigned an enrichment value of -10 and the dropout variants in the clustered double mutant heatmap were arbitrarily assigned an enrichment value of -20. The pheatmap package in R was used to construct the clustered single mutant heatmaps and the NG-CHM[1] package in R was used to construct the clustered double mutant heatmap. Euclidean distances were used for hierarchical clustering of heatmaps.

The appropriate files for DeepLasso are contained in DeepLasso.zip (which contains the Python code for DeepLasso, the training dataset, and the test dataset).

**Expression and Purification of Ubonodin Variants**

pUboABCD variants were transformed into electrocompetent *E. coli* BL21 cells and grown in M9 minimal media supplemented with 40 μg/mL of each canonical amino acid, 0.5 μg/mL thiamine, and 100 μg/mL ampicillin at 37 °C. Once the $OD_{600}$ reached 0.2, cultures were induced with 1 mM IPTG and grown at 20 °C for 20 hours. Cultures were centrifuged at 4,000 g for 20 minutes, and the supernatant was applied to a Thermo Fischer HyperSep 6 mL C8 column. The column was activated with 6 mL of methanol and washed with 12 mL of water prior to applying the supernatant to the column. The column was then washed with 12 mL of water and the extract was eluted with 6 mL of methanol. The methanol was evaporated with a rotary evaporator prior to being resuspended in 1 mL of 75/25 water/acetonitrile per 1 L of culture. Supernatant extracts were injected onto an HPLC using water and acetonitrile with 0.1% trifluoroacetic acid in which 0-1 min ran 10% acetonitrile, 1-20 min ran 10-50% acetonitrile with a linear gradient, and 20-25 min ran 50-90% acetonitrile with a linear gradient. Fractions were collected using the collection windows specified in Table S9 and lyophilized. Injecting these HPLC fractions onto an LC-MS indicated that ubonodin N11W and ubonodin N11W H17T were expressed at levels too low to continue purification.

Ubonodin A7P, ubonodin R12F, ubonodin H17G, ubonodin A7G N11M, ubonodin A7P I16A, and ubonodin R12V H17G were determined to be pure after purifying with the previously stated HPLC gradient. Ubonodin M14N, ubonodin I16D, ubonodin I16E, ubonodin R12F W19G, ubonodin S5T I16D, and ubonodin I16E D23A required a second round of HPLC purification. The lyophilized fractions were resuspended in 75/25 water/acetonitrile and injected onto an HPLC using water and acetonitrile with 0.1% trifluoroacetic acid in which 0-1 min ran 10% acetonitrile, 1-20 min ran 10-28% acetonitrile with a linear gradient, and 20-25 min ran 28-48% acetonitrile with a linear gradient. Ubonodin S5H I16P also required a second round of HPLC purification. The lyophilized fraction was resuspended in 75/25 water/acetonitrile and injected onto an HPLC using water and acetonitrile with 0.1% trifluoroacetic acid in which 0-2 min ran 10% acetonitrile, 2-20 min ran 10-30% acetonitrile with a linear gradient, and 20-25 min ran 30-90% acetonitrile with a linear gradient. Collection windows are specified in Table S9. Purity was verified by injecting pure samples onto an LC-MS, and peptides were resuspended in the solvent appropriate for their hydrophobicity, as specified in Table S9.

**Antimicrobial Assay**

The *Burkholderia* strains used were *B. cenocepacia* AU0756, and *B. cenocepacia* AU24326. The strains were grown at 32 °C. Streaked plates of *B. cenocepacia* AU0756 were grown for 48 hours while streaked plates of *B. cenocepacia* AU24326 were grown for 72 hours.

Broth microdilution assays were conducted following guidelines provided by the Clinical & Laboratory Standards Institute (CLSI). Two to three colonies from a streaked plate were inoculated into 5 mL of LB media and grown for 16 hours. The dense cultures were then sub-cultured at a 1:100 dilution in 5 mL of LB media and grown until the cultures reached the mid-exponential stage ($OD_{600}$ 0.4-0.6). The cultures were then sub-cultured to $OD_{600}$ 0.0005 in cation-adjusted Mueller Hinton broth along with two-fold serial dilutions of ubonodin variants for a final volume of 100 µL in a 96-well plate. Since acetonitrile inhibits *Burkholderia* growth in liquid media, ubonodin variants that were originally suspended in 50/50 water/acetonitrile were lyophilized and resuspended in PBS. This resulted in the concentrated peptide stocks being relatively clear suspensions and were thoroughly mixed before adding to the 96-well plates. The plates were grown at 32 °C for 16 hours while shaking at 250 rpm before measuring the $OD_{600}$.

A

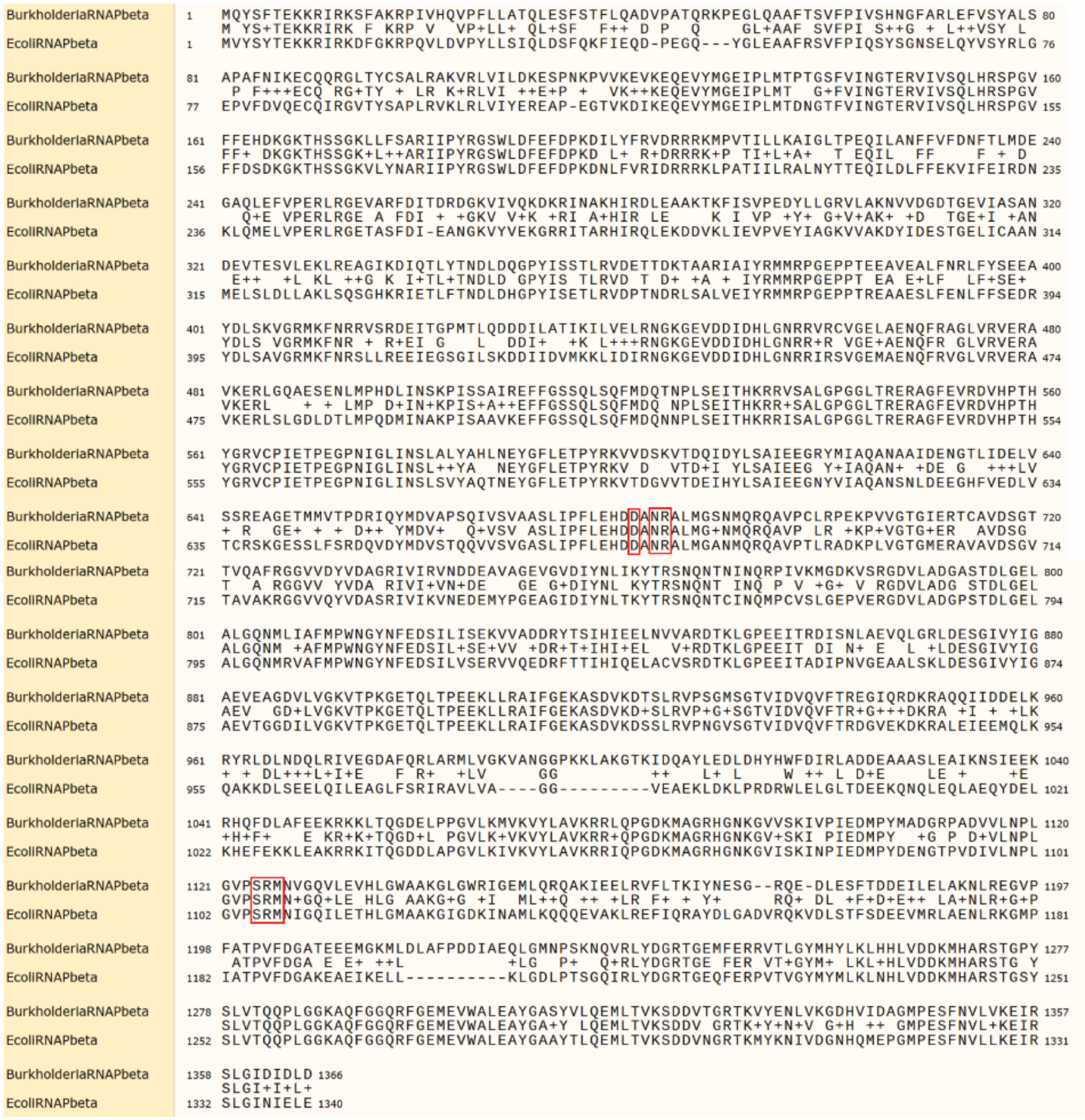


Similarity = 80.28%
Identity = 65.96%

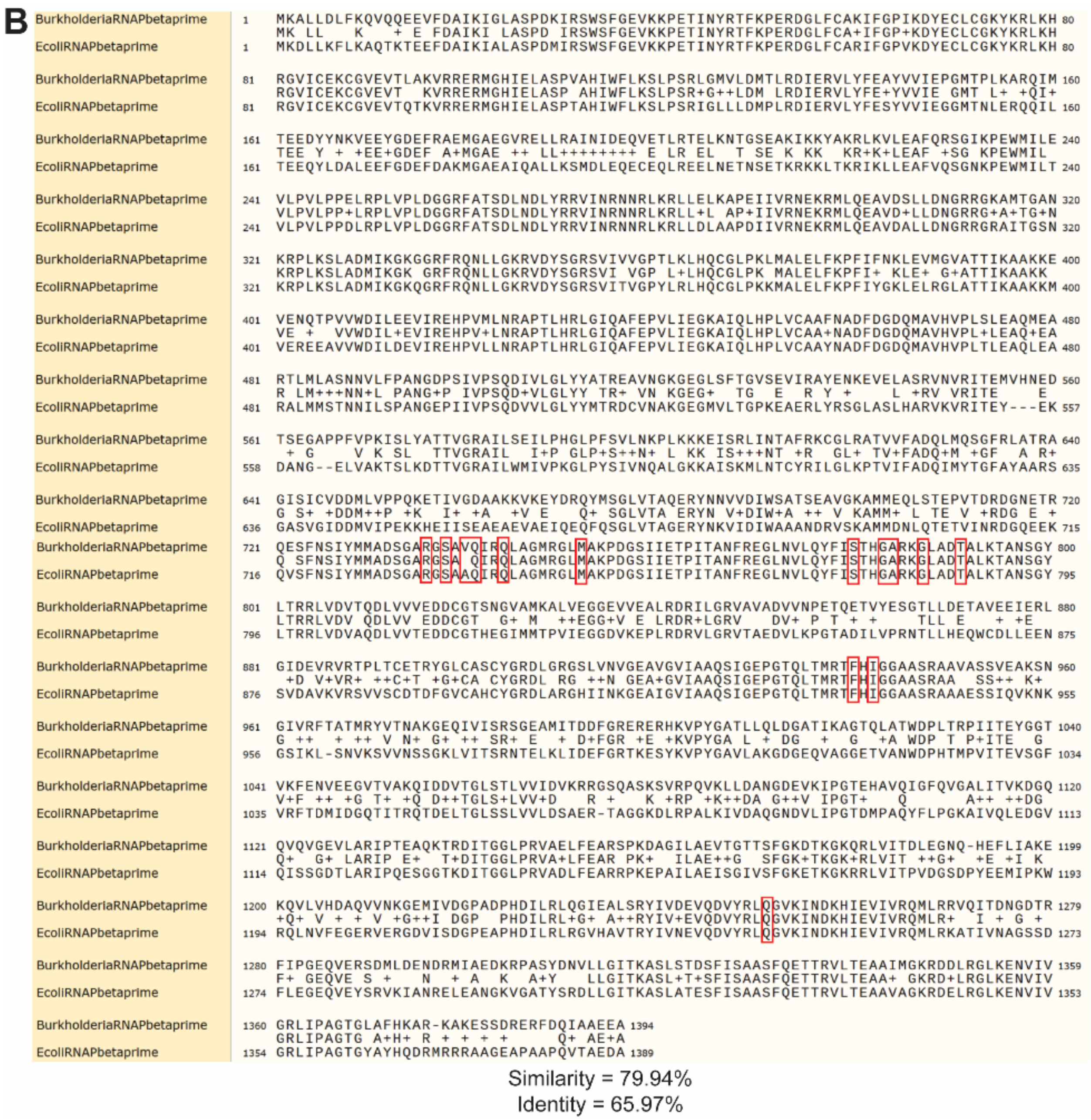


**Figure S1**. Alignment of the *Burkholderia cepacia* RNAP subunits with the *E. coli* RNAP subunits that have been shown to interact with microcin J25,[2] **A**: the β subunit and **B**: the β’ subunit. Residues that directly interact with microcin J25 (D675, N677, R678, S1105, R1106, and M1107 in the β subunit and R731, S733, A735, Q736, Q739, M747, S775, G778, A779, G782, T786, F935, I937, and Q1225 in the β’ subunit) are boxed in red. All these residues are conserved in the *B. cepacia* RNAP subunits except for A735 in the β’ subunit which is substituted to valine instead.

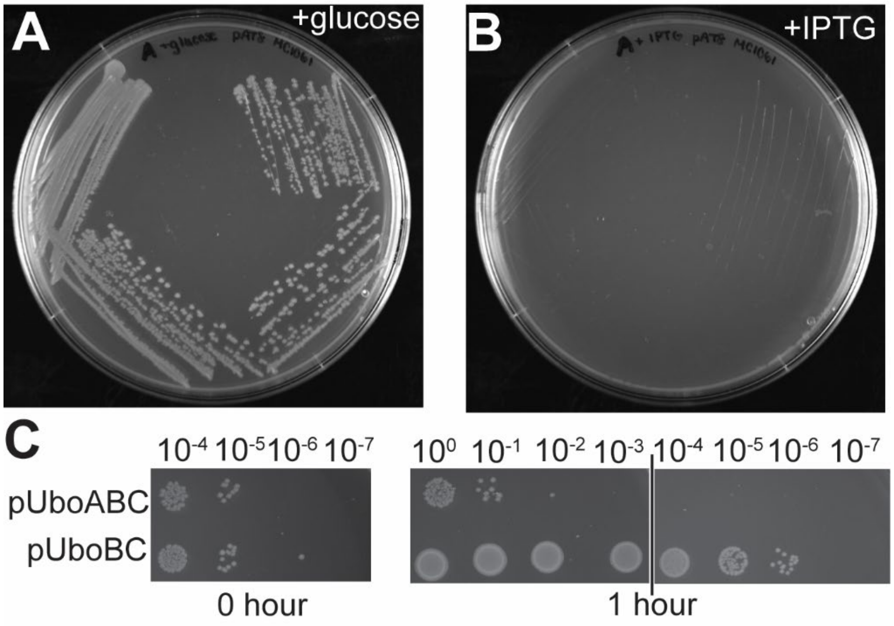


**Figure S2. A**: Streak of *E. coli* pUboABC onto an agar plate supplemented with 100 µg/mL ampicillin (to select for the plasmid) and 0.2% w/v glucose. This condition represses leaky expression of *uboA*, and colonies grew normally. **B**: Streak of *E. coli* pUboABC onto an agar plate supplemented with 100 µg/mL ampicillin and 1 mM IPTG. This condition induces expression of *uboA*, and no colonies grew. **C:** Spot dilution assay of *E. coli* pUboABC and *E. coli* pUboBC. Liquid cultures were plated immediately prior to induction (0 hour spots) and 1 hour after induction (1 hour spots). *E. coli* pUboABC had a clear growth deficiency 1 hour after induction compared to *E. coli* pUboBC.

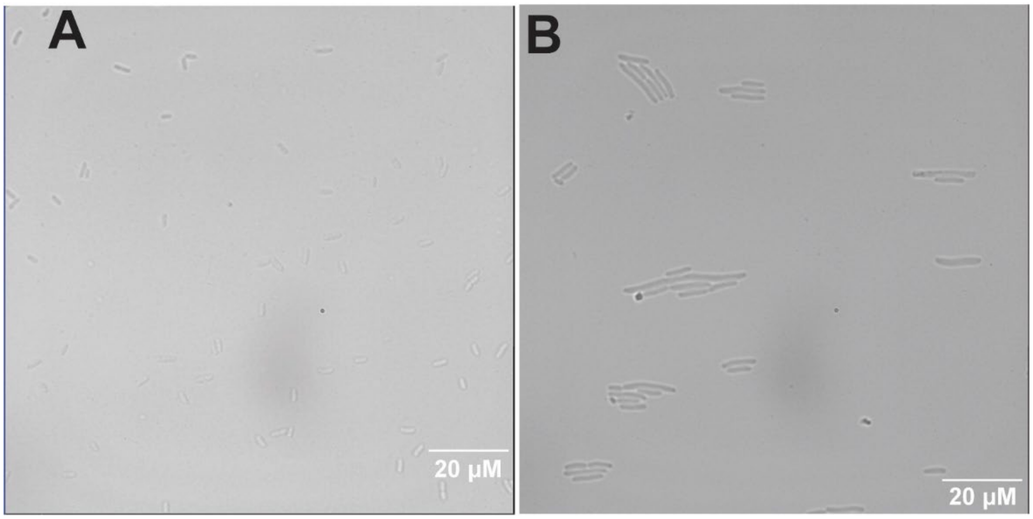


**Figure S3.** Microscopy on *E. coli* pUboBC and *E. coli* pUboABC. **A**: *E. coli* pUboBC cells had a normal rod shape two hours after induction with 100 μM IPTG. **B**: *E. coli* pUboABC cells two hours after induction with 100 μM IPTG tended to be longer than the cells depicted in panel A, indicative of filamentation.

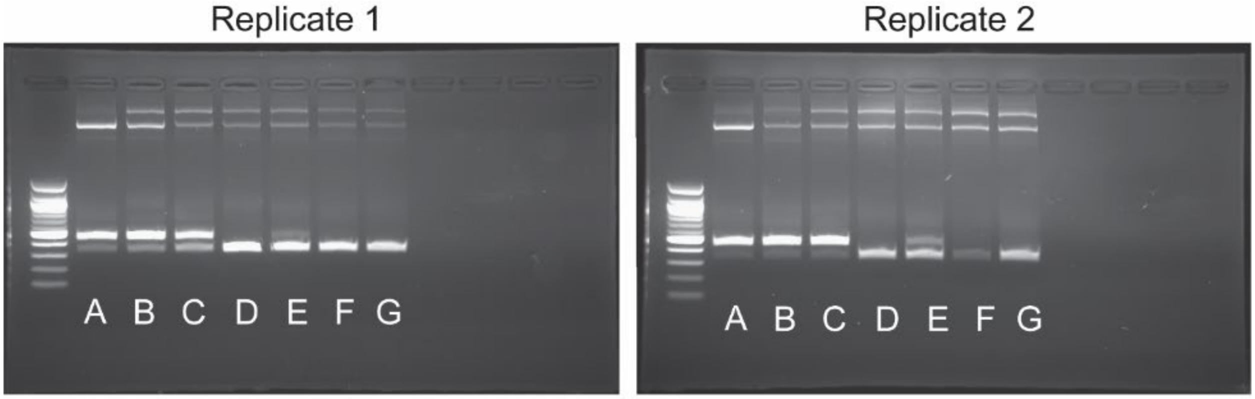


| PCR | DNA Template |
|---|---|
| A | 90% pUboABC, 10% pUboBC |
| B | Miniprepped DNA from transforming into *E. coli* MC1061 |
| C | Miniprepped DNA from co-culture, immediately prior to IPTG induction |
| D | Miniprepped DNA from co-culture, 1 hour after 100 µM IPTG induction |
| E | Miniprepped DNA from co-culture, 1 hour after 10 µM IPTG induction |
| F | Miniprepped DNA from co-culture, 2 hours after 10 µM IPTG induction |
| G | Miniprepped DNA from co-culture, 3 hours after 10 µM IPTG induction |

**Figure S4**. DNA that was PCR-amplified from co-cultures of *E. coli* pUboABC and *E. coli* pUboBC. The longer amplicon (447 bp band) corresponds to pUboABC and the shorter amplicon (336 bp band) corresponds to pUboBC. The co-culture primarily consisted of *E. coli* pUboABC until induction, after which the *E. coli* pUboBC cells began to overtake the co-culture.

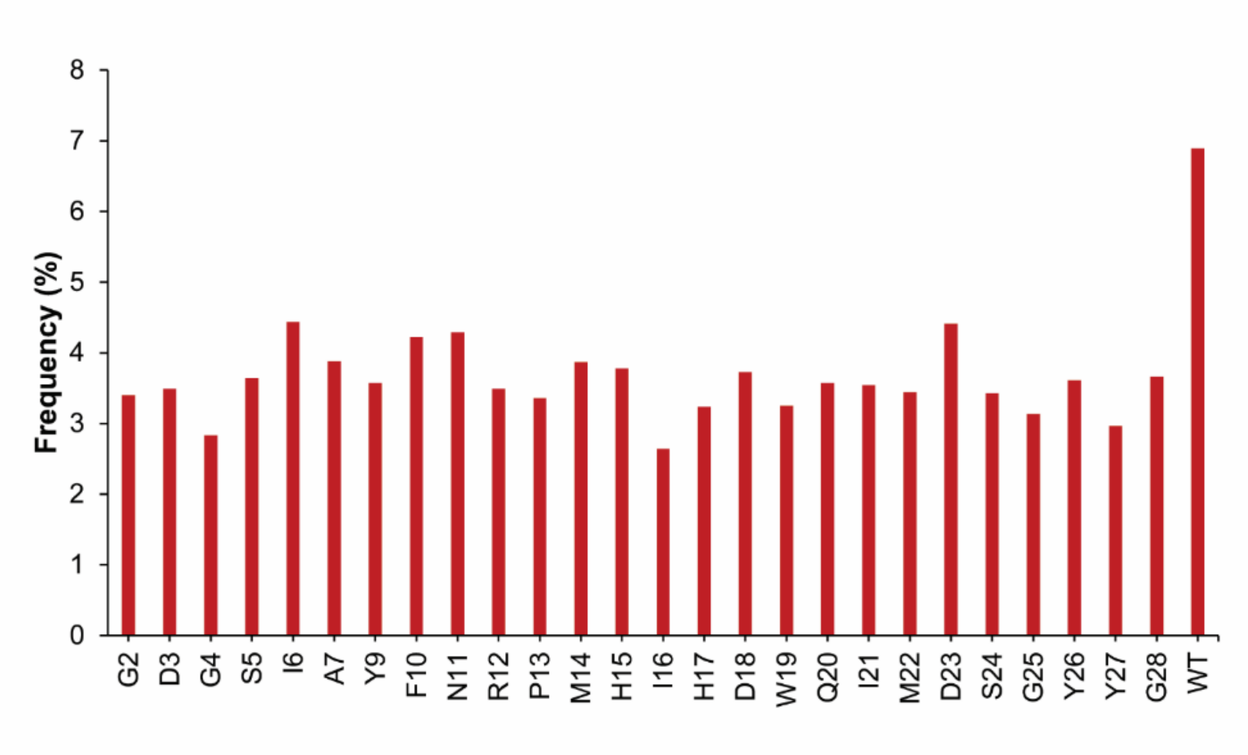


**Figure S5**. The frequencies of point variants in the cloning transformation library of the single mutant library were summed for each residue after sequencing on the NovaSeq (Equation S6), revealing that the residue-specific libraries were combined in roughly equimolar amounts. WT ubonodin was present at higher levels than expected, which could be due to the WT bias when conducting PCR with NNK codons, which has been documented previously.[3]

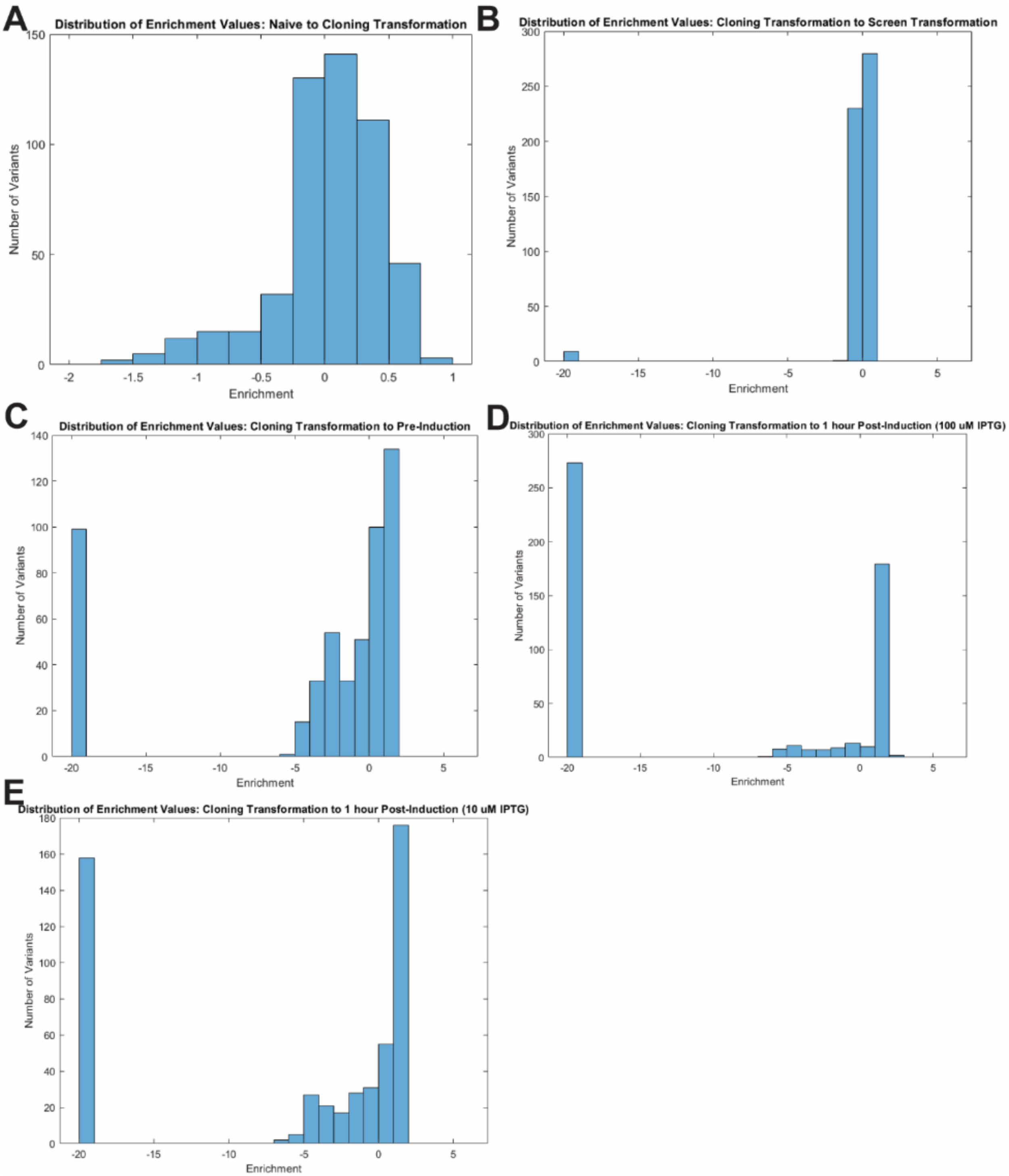


**Figure S6.** Histograms depicting the distribution of point variants' enrichment values at different steps of the single mutant screen (data from NovaSeq single mutant screen). Dropout variants were arbitrarily assigned an enrichment value of -20. Comparisons are between **A:** The naïve library and the cloning transformation library. Enrichment values have a small range, with 80% of variants falling between an enrichment value of -0.5 and 0.5. Only two variants (D3F and D3M) were not present in the cloning transformation library, likely because they had a low read count in the naïve library (626 reads and 554 reads respectively). **B:** The cloning transformation library and the screen transformation sample. **C**: The cloning transformation library and the pre-IPTG sample. **D**: The cloning transformation library and the 1-hour post-100 μM IPTG induction sample. **E:** The cloning transformation library and the 1-hour post-10 μM IPTG induction sample.

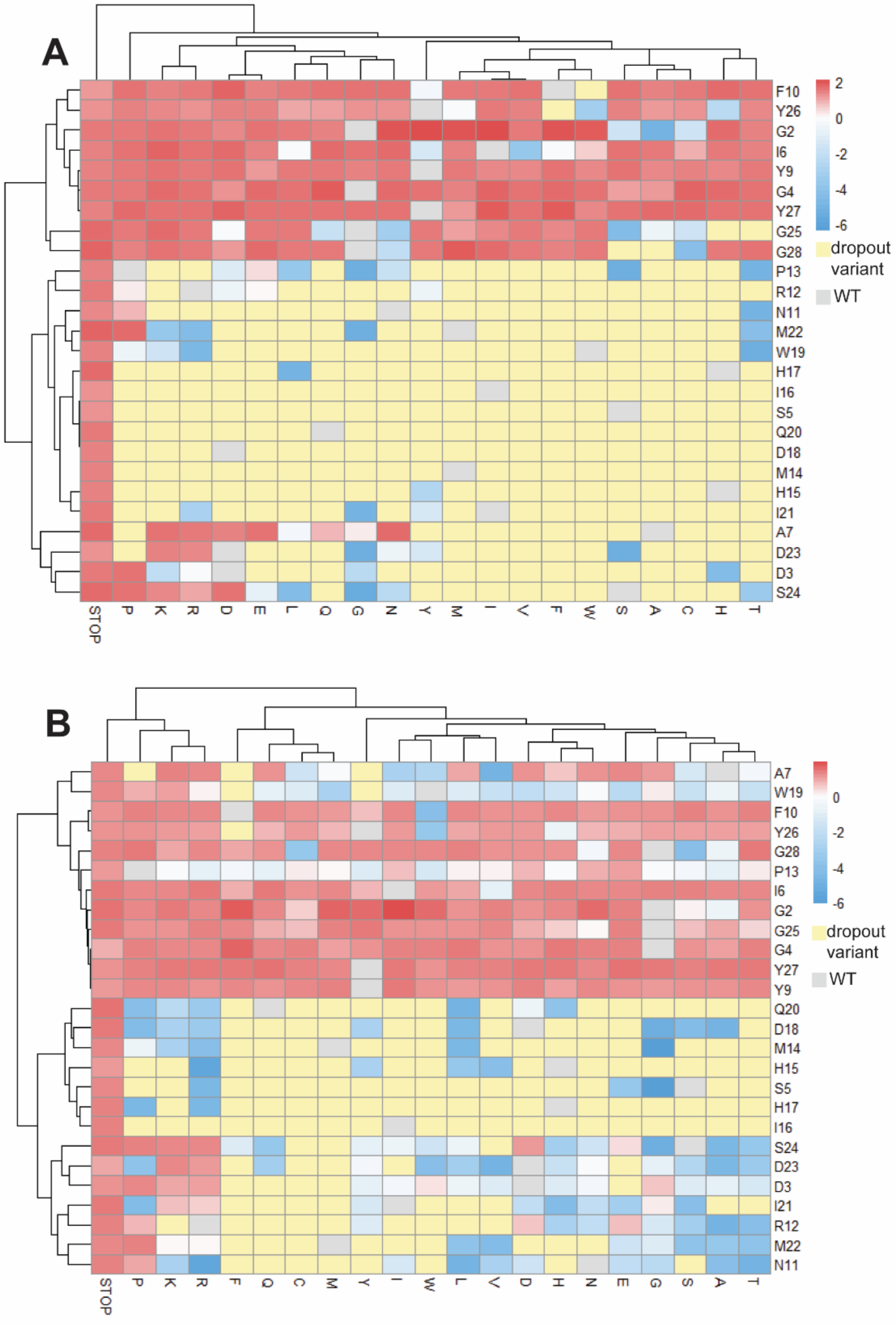


**Figure S7**. Hierarchical clustering of enrichment values for the single mutant library after induction with **A**: 100 μM IPTG or **B**: 10 μM IPTG. Data is obtained from the NovaSeq sequencing run of the single mutant library.

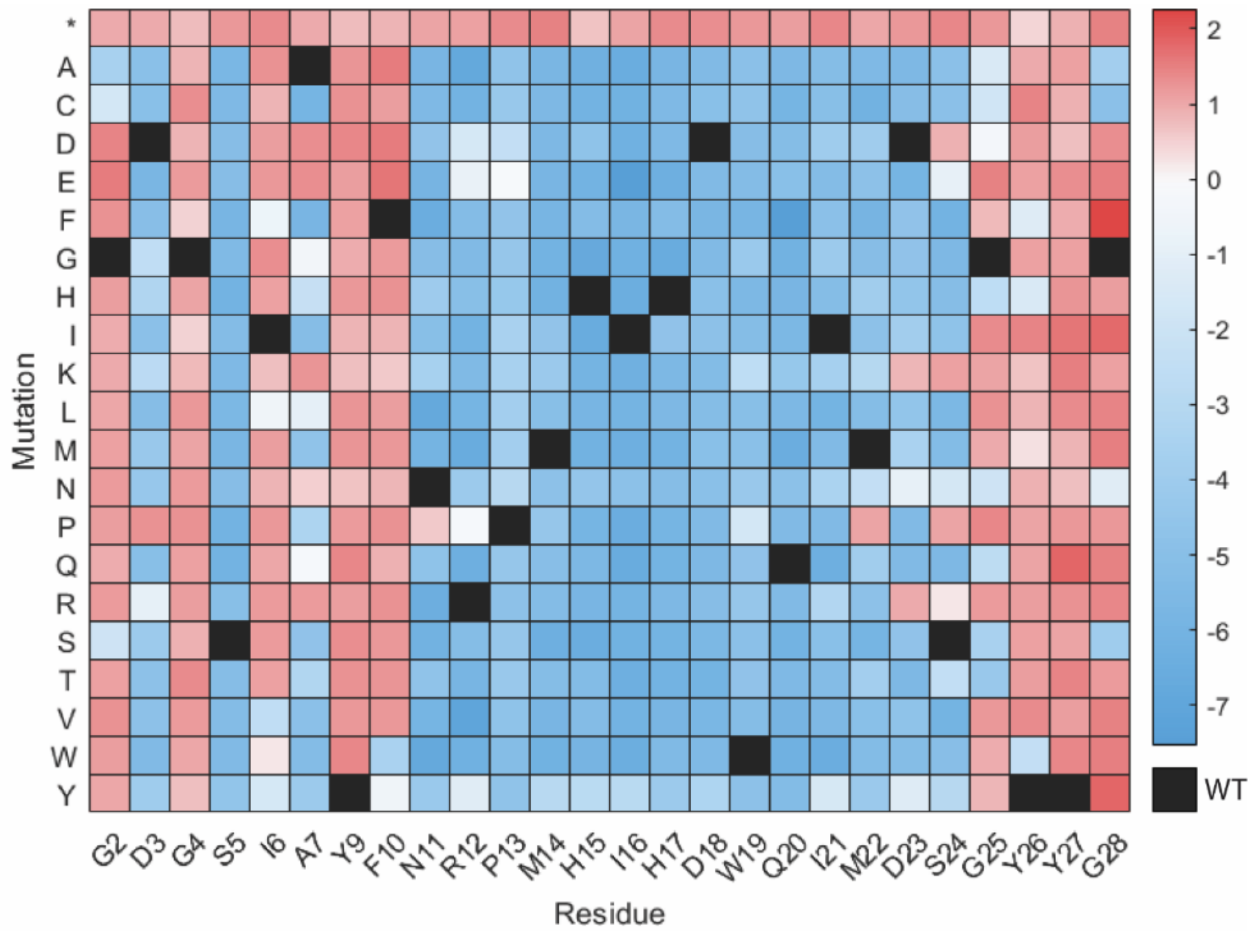


**Figure S8**. Enrichment values of the point variants present in the double mutant library (NovaSeq sequencing run on the double mutant library) after induction with 100 μM IPTG. The heatmap displays the same pattern of RNAP-inhibiting and non-RNAP inhibiting point variants as was observed in the single mutant library; compare to Figure 3A in the main text from the NovaSeq sequencing run on the single mutant library. There are no dropout single aa variants in the double mutant library, which is likely because we used more reads to sequence the double mutant library than the single mutant library.

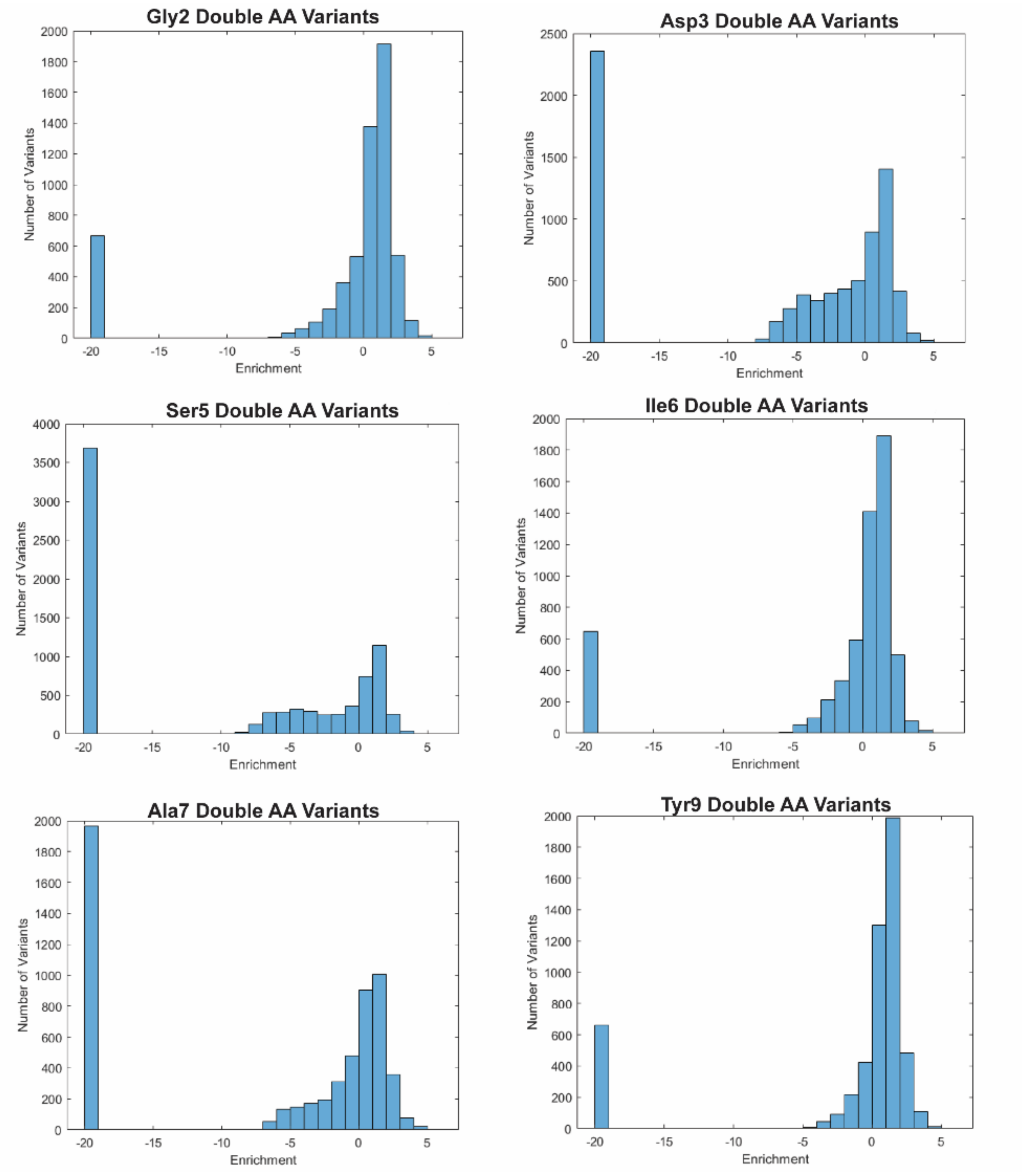
Gly2 Double AA Variants
Number of Variants
Enrichment
Asp3 Double AA Variants
Number of Variants
Enrichment
Ser5 Double AA Variants
Number of Variants
Enrichment
Ile6 Double AA Variants
Number of Variants
Enrichment
Ala7 Double AA Variants
Number of Variants
Enrichment
Tyr9 Double AA Variants
Number of Variants
Enrichment

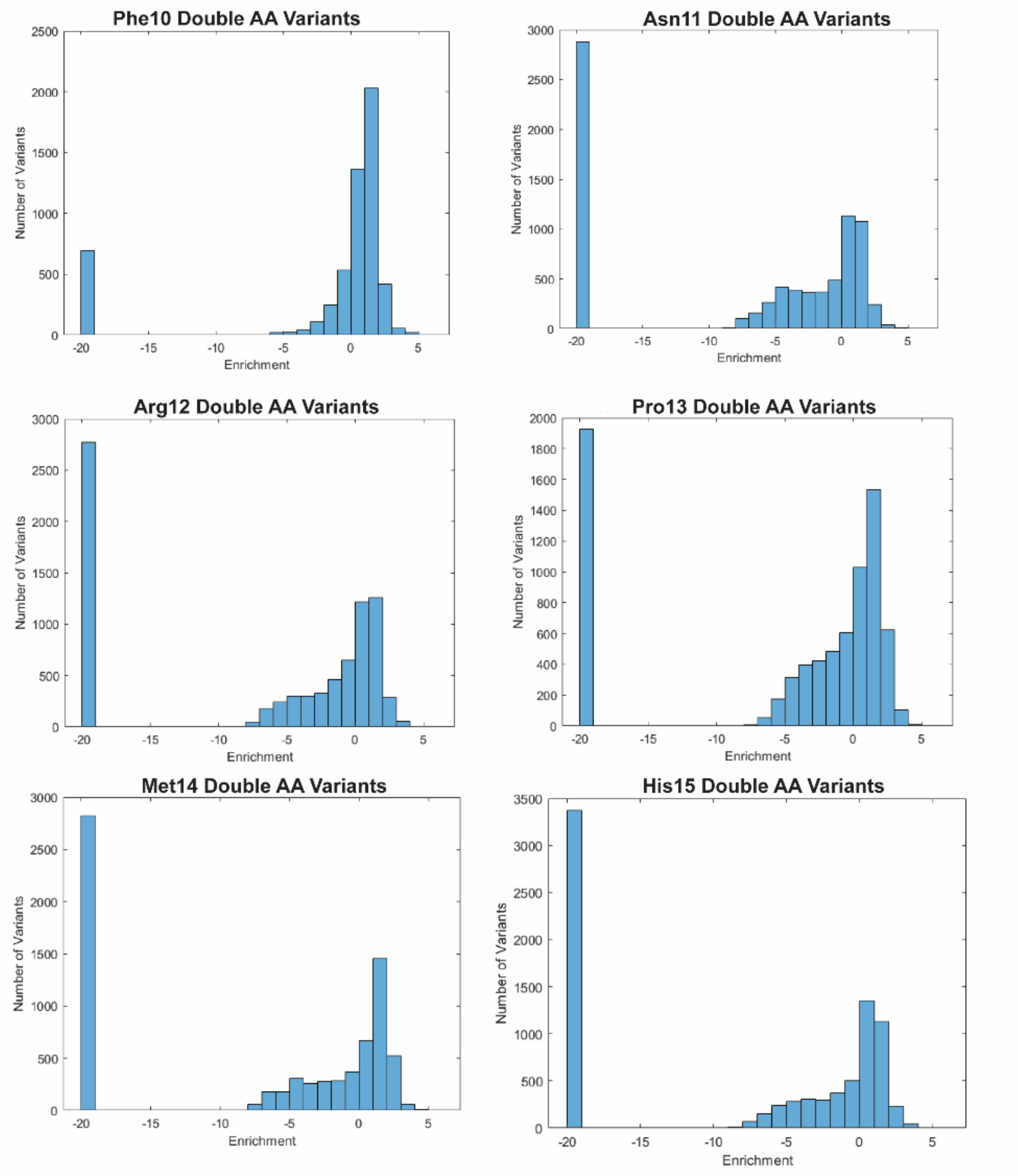

Phe10 Double AA Variants
Asn11 Double AA Variants
Arg12 Double AA Variants
Pro13 Double AA Variants
Met14 Double AA Variants
His15 Double AA Variants
Number of Variants
Enrichment

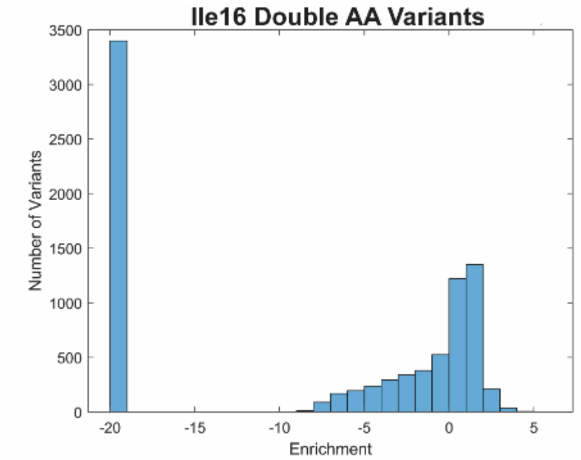
Ile16 Double AA Variants
Number of Variants
3500
3000
2500
2000
1500
1000
500
0
-20
-15
-10
-5
0
5
Enrichment

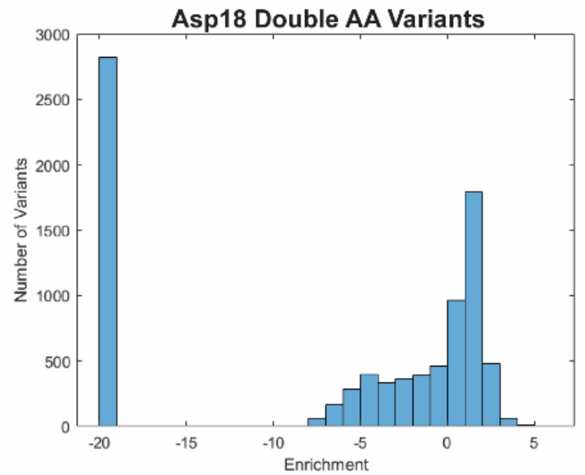
Asp18 Double AA Variants
Number of Variants
3000
2500
2000
1500
1000
500
0
-20
-15
-10
-5
0
5
Enrichment

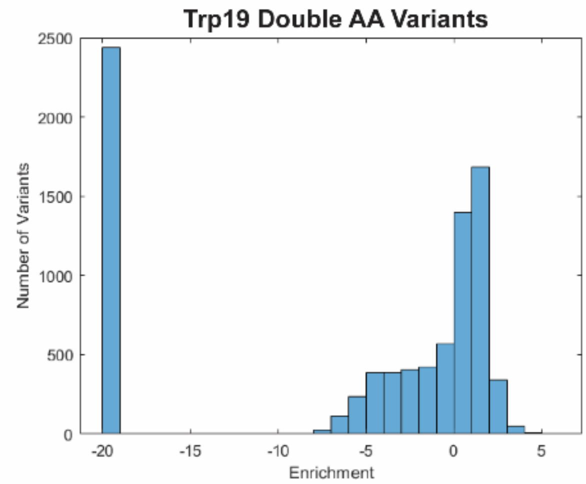
Trp19 Double AA Variants
Number of Variants
2500
2000
1500
1000
500
0
-20
-15
-10
-5
0
5
Enrichment

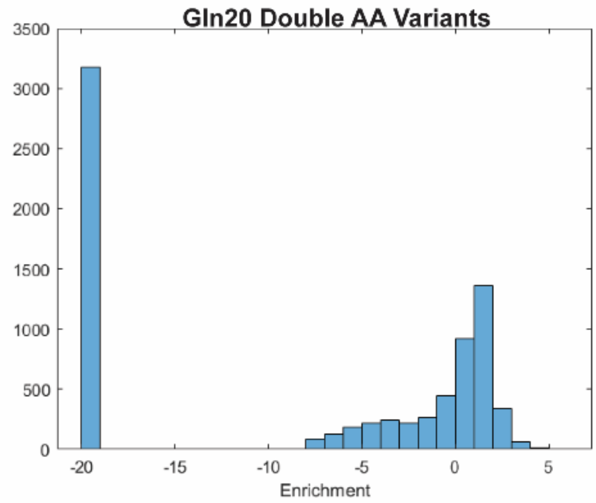
Gln20 Double AA Variants
3500
3000
2500
2000
1500
1000
500
0
-20
-15
-10
-5
0
5
Enrichment

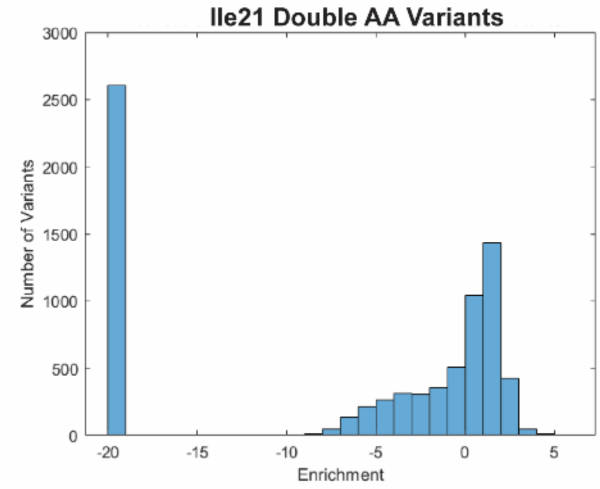
Ile21 Double AA Variants
Number of Variants
3000
2500
2000
1500
1000
500
0
-20
-15
-10
-5
0
5
Enrichment

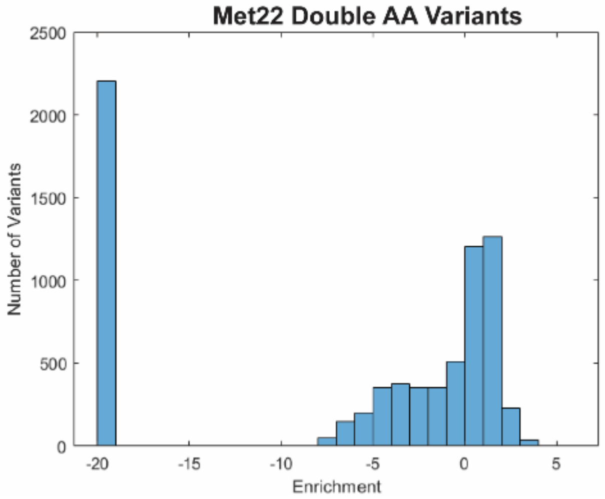
Met22 Double AA Variants
Number of Variants
2500
2000
1500
1000
500
0
-20
-15
-10
-5
0
5
Enrichment

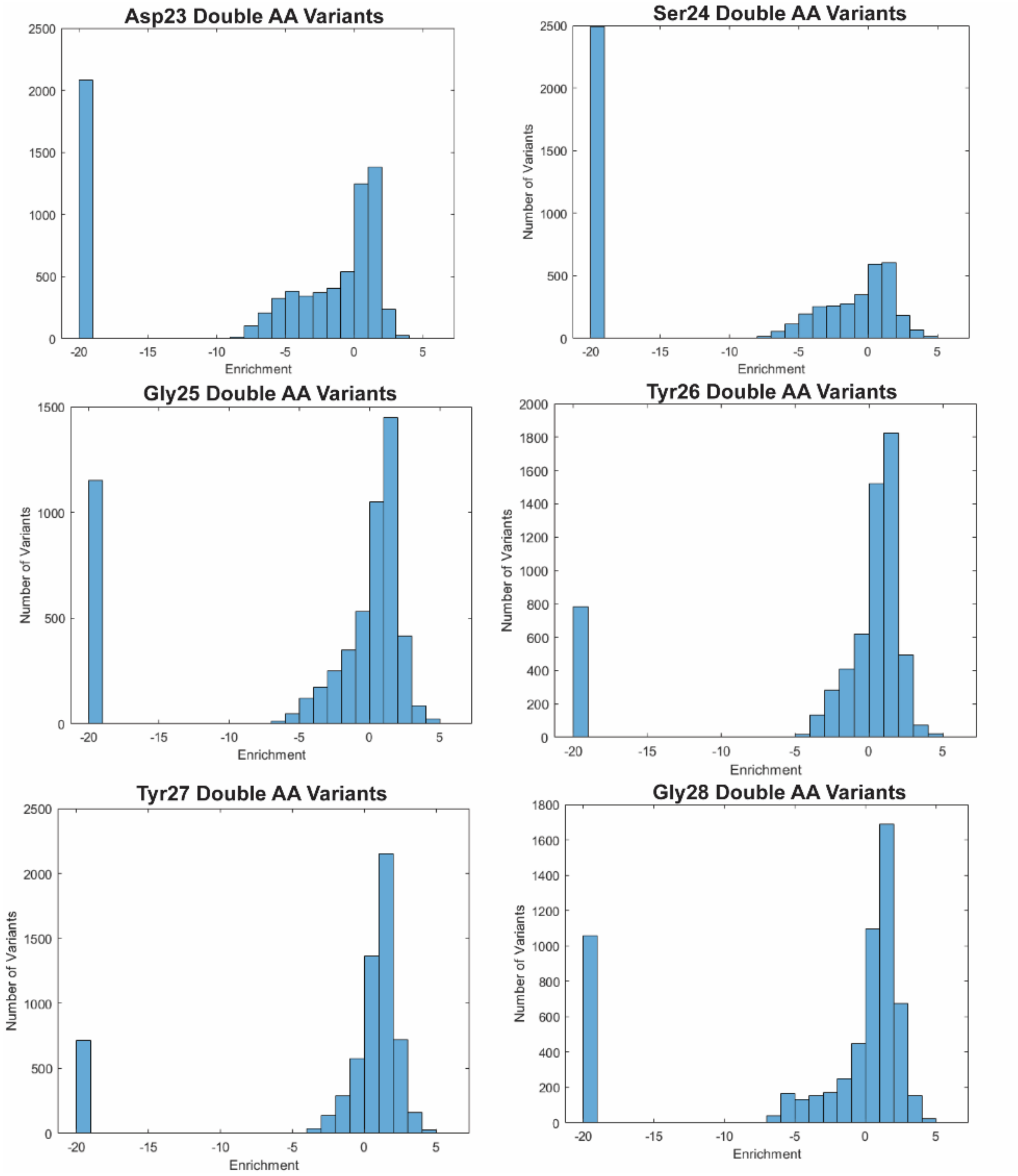


**Figure S9**. Histograms displaying the spread of enrichment values for each residue-specific subset of the double mutant library, excluding the Gly4 and His17 histograms (shown in Figure 5A-B) and positions that were not mutated. Dropout variants were arbitrarily assigned an enrichment value of -20.

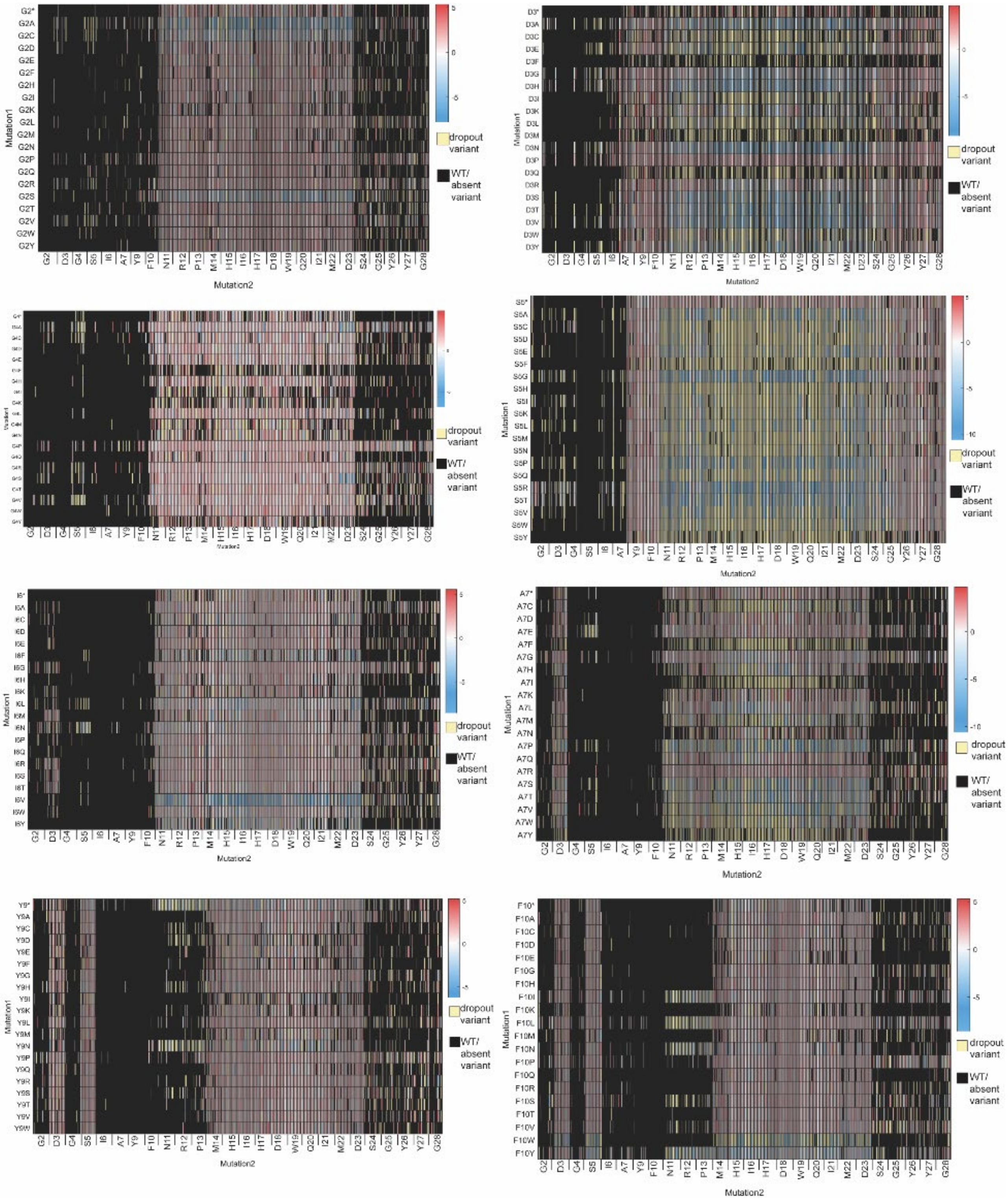
Mutation1
Mutation2
G2 D3 G4 S5 I6 A7 Y9 F10 N11 R12 P13 M14 H15 I16 H17 D18 W19 Q20 I21 M22 D23 S24 G25 Y26 Y27 G28
dropout variant
WT/ absent variant
5
0
-5
-10

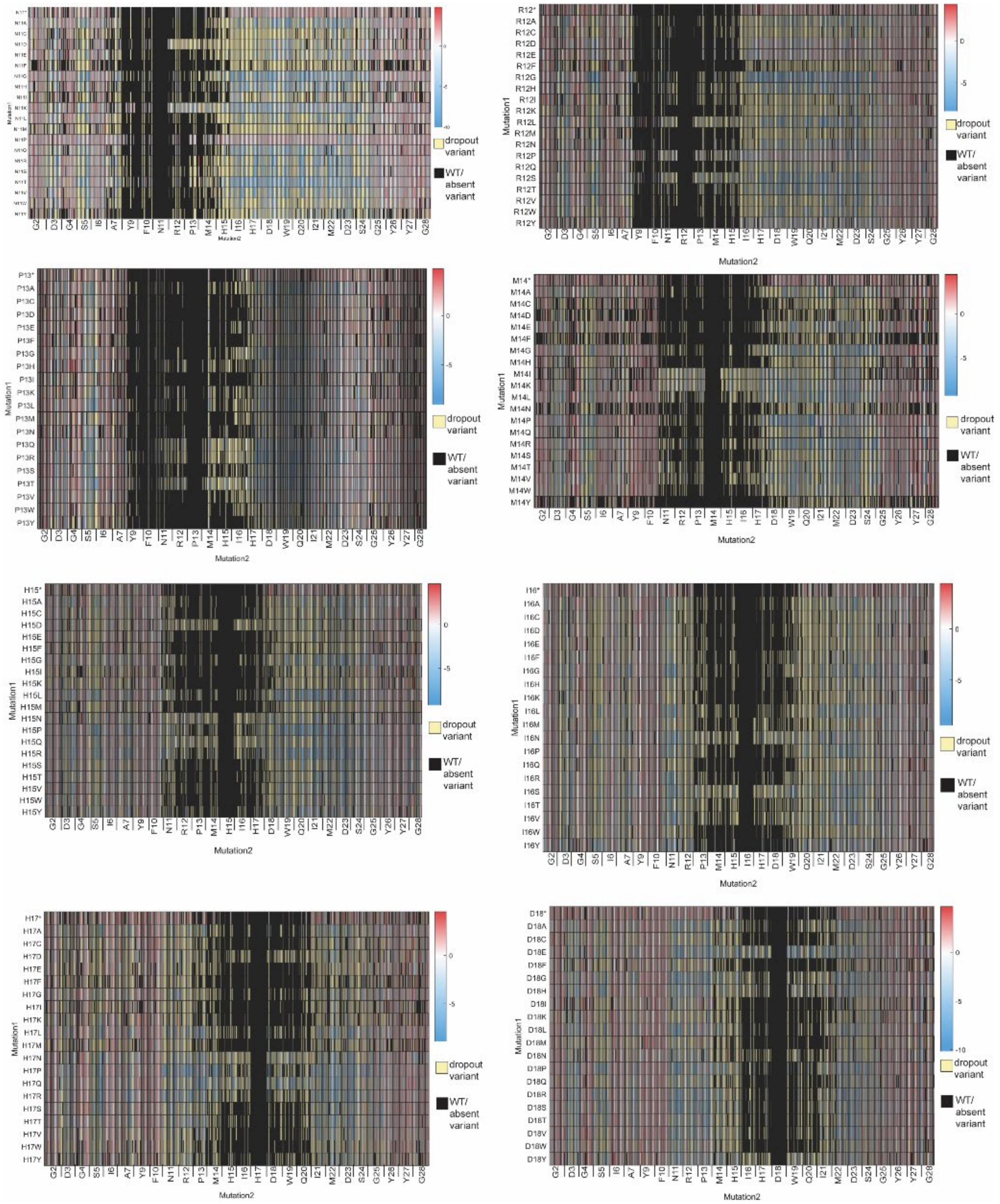
Mutation1
Mutation2
dropout variant
WT/ absent variant

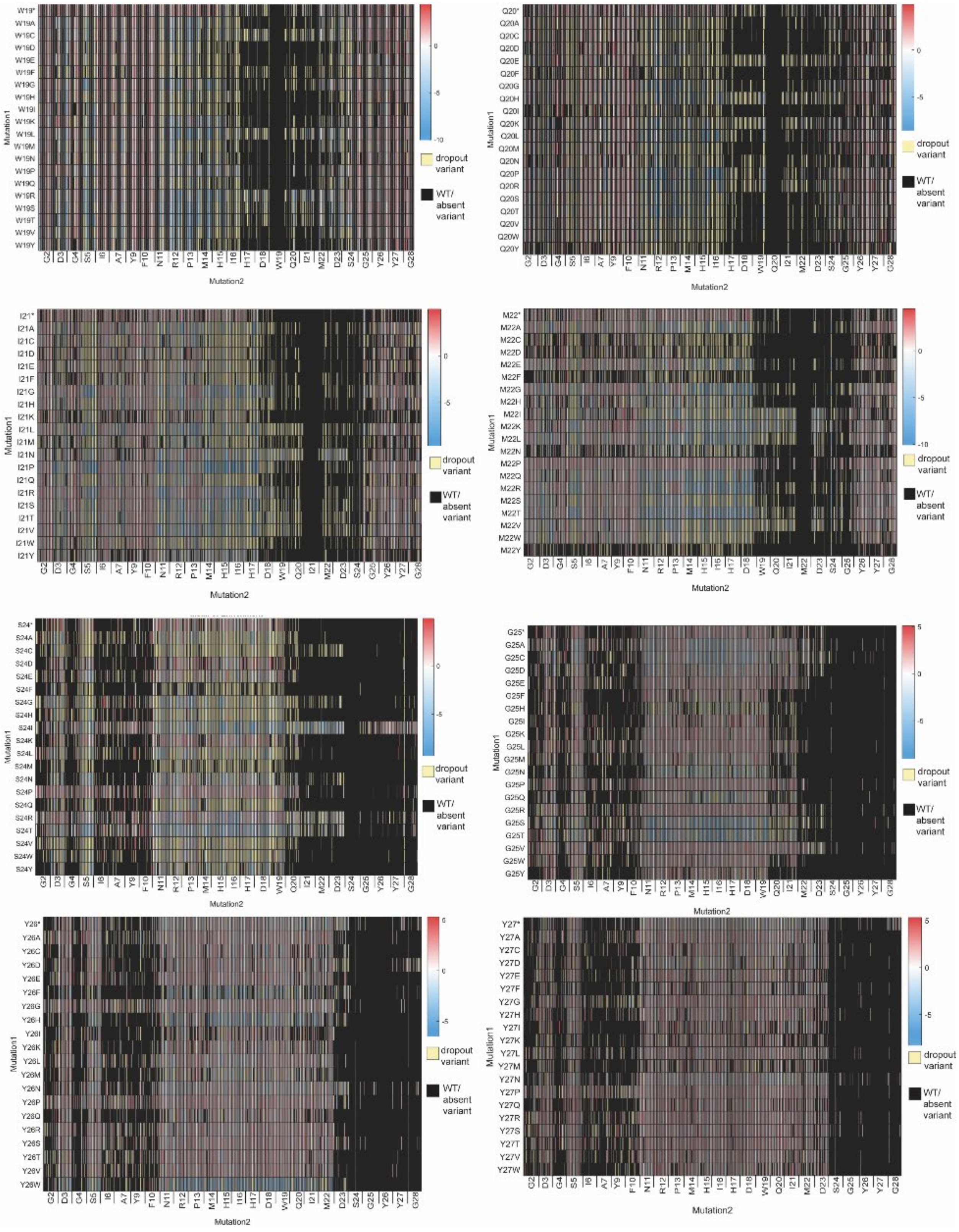
Mutation1
Mutation2
dropout variant
WT/ absent variant

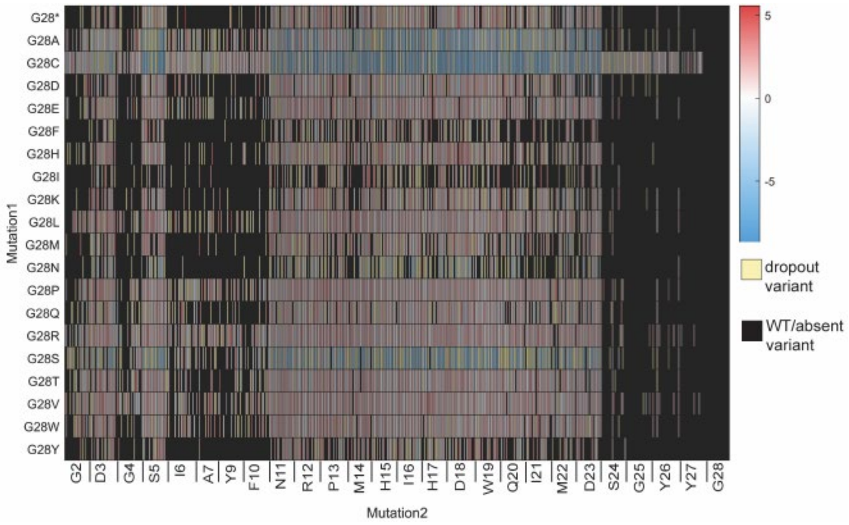


**Figure S10**. Non-clustered heatmaps of each residue-specific subset in the double mutant library.

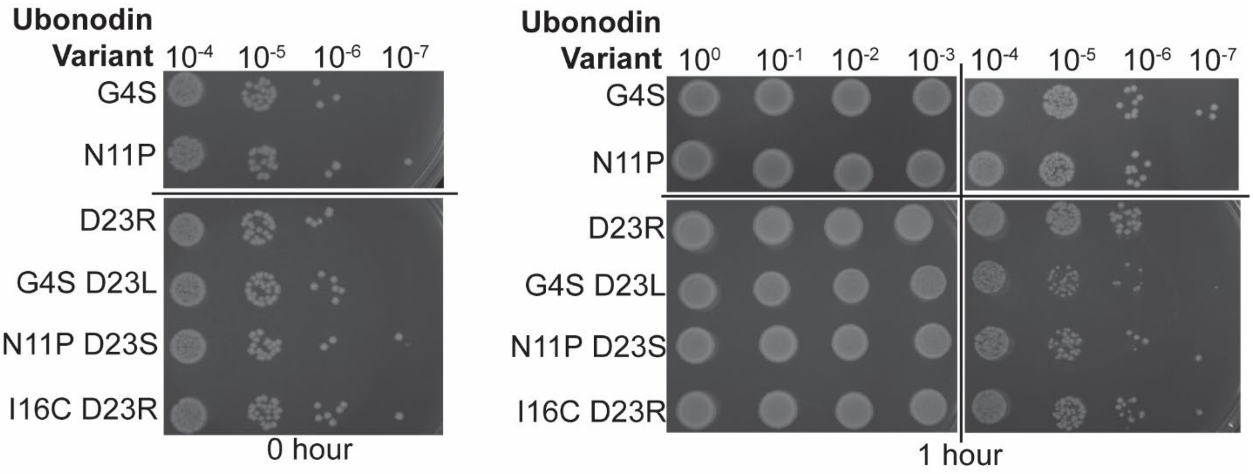


**Figure S11.** Spot dilution assay of the rescue variants and point variants without the compensatory mutation (in a pUboABC backbone). Liquid cultures were plated immediately prior to induction (0 hour spots) and 1 hour after induction (1 hour spots). The spots of point variants have healthy, normal-sized colonies 1 hour after induction whereas the spots of rescue variants have small, sick colonies 1 hour after induction. Biological duplicates of this assay were conducted, and this replicate is representative of the results from both replicates.

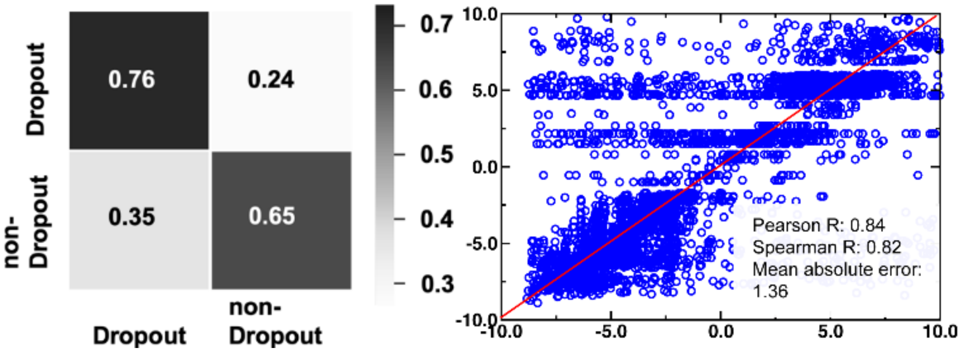


**Figure S12.** (*Left*) Confusion matrix for the classifier of DeepLasso in the training set. The matrix shows binary classification of dropout versus non-dropout variants with predicted outcomes on the x-axis and experimental observation on the y-axis. Grayscale is used to represent the magnitude of probability (i.e., high: black; low: white). (*Right*) Regression analysis for the regressor of DeepLasso in the training set. The linear correlation between experimental vs. predicted enrichment values is shown along with Pearson correlation coefficient, Spearman correlation coefficient, and mean absolute error.

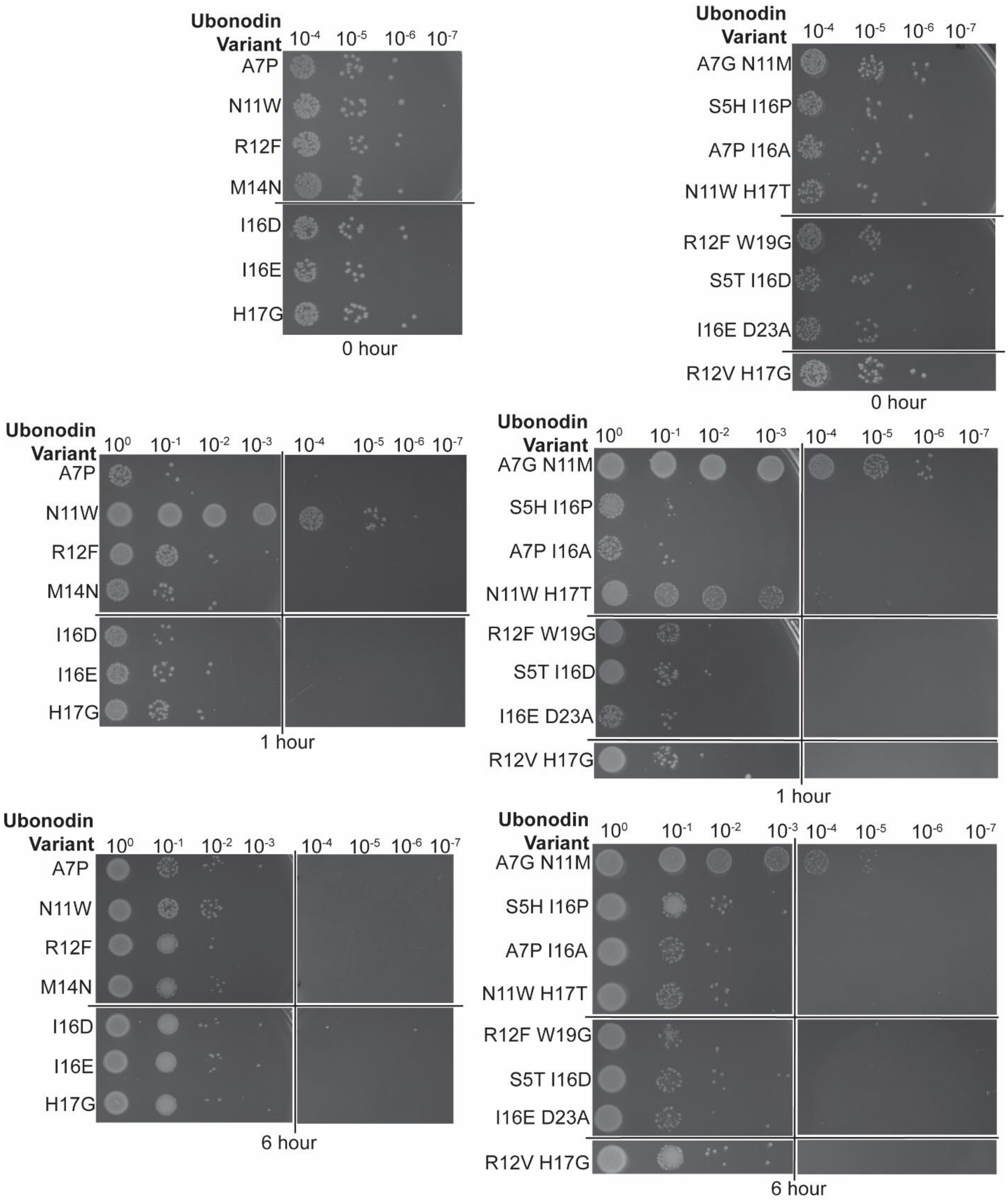


**Figure S13**. Spot dilution assays for the ubonodin variants (in a pUboABC backbone) identified from the screen with potential activity. The toxic effect of most variants was observed 1 hour after induction, though the toxic effect of ubonodin N11W and ubonodin A7G N11M was only observed 6 hours after induction. The apparent differences in toxicity could be because the screen incorporated information on leaky expression prior to induction, which cannot be done with the spot dilution assay. Furthermore, heterologous expression of ubonodin N11W in a pUboABCD vector demonstrated that the variant is expressed at very low levels, so it may take more time for ubonodin N11W to accumulate to levels that cause a visible difference in the spot dilution assay. Biological duplicates of this assay were conducted, and the replicate shown is representative of both duplicates.

**Table S1.** Enrichment values of rescue variants.

| Single Amino Acid Variant | Single Amino Acid Variant Enrichment | Residue Implicated with Many Rescue Variants | Rescue Variant Enrichment | |
|---|---|---|---|---|
| **G4S** | 1.1 | Asp23 | G4S D23A: -4.5<br>G4S D23C: -3.4<br>G4S D23E: -1.0<br>G4S D23F: dropout variant<br>G4S D23G: -2.2<br>G4S D23L: -4.926<br>G4S D23M: -3.2 | G4S D23N: -0.1<br>G4S D23P: -2.9<br>G4S D23Q: -0.4<br>G4S D23S: -3.5<br>G4S D23V: -4.6<br>G4S D23W: -0.9 |
| **N11P** | 0.97 | Asp23 | N11P D23A: -5.1<br>N11P D23C: -5.6<br>N11P D23F: dropout variant<br>N11P D23G: -1.3<br>N11P D23I: dropout variant<br>N11P D23L: -4.6<br>N11P D23M: -5.0<br>N11P D23N: -0.3 | N11P D23P: -5.4<br>N11P D23Q: -2.2<br>N11P D23S: -3.6<br>N11P D23T: -3.3<br>N11P D23V: -4.5<br>N11P D23W: -0.9<br>N11P D23Y: -1.1 |
| **D23K** | 1.2 | Ile16 | I16A D23K: -0.5<br>I16C D23K: -0.3<br>I16D D23K: -2.4<br>I16E D23K: -2.8<br>I16G D23K: -3.8<br>I16H D23K: -0.3<br>I16K D23K: -1.9 | I16N D23K: -1.6<br>I16Q D23K: dropout variant<br>I16R D23K: -0.8<br>I16S D23K: -0.8<br>I16T D23K: -1.7<br>I16V D23K: -0.3<br>I16W D23K: -0.1 |
| **D23R** | 1.2 | Ile16 | I16A D23R: -1.7<br>I16C D23R: -2.9<br>I16D D23R: -3.2<br>I16E D23R: -1.9<br>I16G D23R: -3.1<br>I16H D23R: -1.9<br>I16K D23R: -2.5 | I16M D23R: -0.4<br>I16N D23R: -1.6<br>I16P D23R: -0.1<br>I16Q D23R: -2.1<br>I16R D23R: -2.6<br>I16S D23R: -1.3<br>I16T D23R: -1.8 |
| **Y26L** | 1.1 | Ile16 | I16A Y26L: -0.4<br>I16C Y26L: -0.4<br>I16D Y26L: -1.0<br>I16E Y26L: -0.4<br>I16F Y26L: -0.1<br>I16G Y26L: -1.6<br>I16H Y26L: -0.2<br>I16K Y26L: -1.1 | I16M Y26L: -0.3<br>I16N Y26L: -0.3<br>I16P Y26L: -0.5<br>I16Q Y26L: -1.5<br>I16R Y26L: -0.6<br>I16S Y26L: -0.9<br>I16T Y26L: -0.2 |

**Table S2**. Hyperparameters used in the encoders, classifier, and regressor of DeepLasso.

| | Sequence encoder | Topology encoder | Classifier | Regressor |
|---|---|---|---|---|
| Convolutional Layers | 3 | 3 | 1 | 0 |
| LSTM Layers | 2 | 0 | 0 | 0 |
| Residual Blocks | 0 | 0 | 2 | 1 |
| MLP Layers | 0 | 0 | 1 | 2 |
| Output Channels | 2048 | 768 | N/A | N/A |
| Batch Size | 512 | 512 | 64 | 64 |
| Dropout | 0.5 | 0.5 | 0.5 | 0.5 |
| Learning Rate | 1e-4 | 1e-4 | 2e-4 | 2e-4 |

**Table S3**. Statistics of types of variants involved in the training and test sets for classification and regression.

| | Classifier training | Classifier testing | Regressor training | Regressor testing |
|---|---|---|---|---|
| Wild type | 1 | 0 | 1 | 0 |
| Single aa variant | 370 | 8 | 222 | 19 |
| Double aa variant | 11940 | 997 | 6131 | 581 |
| Triple aa variant | 40875 | 3893 | 1639 | 174 |
| Quadruple aa variant | 2956 | 798 | 103 | 11 |
| Higher-order aa variant | 796 | 85 | 4 | 0 |
| Stop codon | 4745 | 387 | 2230 | 248 |

**Table S4**. Prediction results for single and double aa variants whose MICs were tested. The column variable, *Experimental observation*, refers to the enrichment for a lasso peptide variant observed in the screen. *DeepLasso prediction (classifier)* refers to the prediction outcome of DeepLasso in classifying a mutant to be dropout mutant or non-dropout variant. *DeepLasso prediction (regressor)* refers to enrichment value predicted for non-dropout variants. *Test or training (classifier)* and *Test or training (regressor)* refer to whether the tested variant appears in the training, testing set, or neither.

| Variants | Experimental observation | DeepLasso prediction (classifier) | DeepLasso prediction (regressor) | Test or training (classifier) | Test or training (regressor) |
|---|---|---|---|---|---|
| A7G N11M | Dropout | Dropout | – | Testing | – |
| S5H I16P | Dropout | Dropout | – | Testing | – |
| N11W H17T | Dropout | Non-Dropout | -7.4 | Training | – |
| R12V H17G | Dropout | Dropout | – | Training | – |
| I16E D23A | Dropout | Dropout | – | Training | – |
| S5T I16D | Dropout | Non-Dropout | -5.4 | Testing | – |
| A7P I16A | Dropout | Non-Dropout | -5.3 | Training | – |
| R12F W19G | Dropout | Dropout | – | Training | – |
| H17G | -6.5 | Non-Dropout | -6.1 | Training | Training |
| I16E | -7.5 | Non-Dropout | -4.8 | Training | Training |
| M14N | -4.7 | Non-Dropout | -4.1 | Training | Training |
| R12F | -5.3 | Dropout | – | Training | Training |
| A7P | -3.4 | Non-Dropout | -3.2 | Training | Training |
| I16D | -6.2 | Non-Dropout | -7.8 | Training | Training |
| N11W | Dropout | Dropout | – | Training | – |

**Table S5**. MIC values of various antibiotics against the *B. cenocepacia* clinical isolates used in this study.[4-6] WT ubonodin has a lower MIC than the other antibiotics reported in the literature. However, it is important to note that while we used the CLSI broth microdilution method to measure the MIC of ubonodin, the CLSI agar dilution method was used in the literature to measure the MIC of the other compounds. Two values were given for combination treatments: an MIC of 32/4 for ceftazidime-avibactam means that when the two antibiotics are used together, the bacteria must be treated with 32 μg/mL ceftazidime and 4 μg/mL avibactam to inhibit cell growth.

| Compound | ***B. cenocepacia*** **AU0756 MIC (μg/mL)** | ***B. cenocepacia*** **AU24326 MIC (μg/mL)** |
|---|---|---|
| **WT ubonodin** | 0.8 – 1.6 | 6.4 |
| **Tobramycin** | 512 | 512 |
| **Ciprofloxacin** | 256 | 8 |
| **Minocycline** | 8 | 32 |
| **Imipenem** | 128 - 256 | 128 |
| **Ceftazidime** | 128 | 32 |
| **Trimethoprim-sulfamethoxazole** | 8/152 | 4/76 |
| **Ceftazidime-avibactam** | 32/4 | 8/4 |
| **Imipenem-relebactam** | 32/4 | Not reported |
| **Piperacillin-avibactam** | 4/4 | Not reported |
| **Piperacillin** | Not reported | 512 |

**Table S6**. Primers used in this study.

| Primer Name | Sequence |
|---|---|
| pQE-80 *Nhe*I For | GATCGCTAGCCATCAATTAAGA |
| pAT8 *Nco*I Rev | GCATCCATGGTCACTTCATGGTCTGGTTAGTTC |
| *Xho*I For Lib | AGCCGTCTTCACCTCGAG |
| *Nhe*I Rev Lib | TCTTAATTGATGGCTAGCTTG |
| G2 For Lib | GACAATGGGANNKGATGGCAGCA |
| G2 Rev Lib | TGCTGCCATCMNNTCCCATTGTC |
| D3 For Lib | GAGGCNNKGGCAGCAT |
| D3 Rev Lib | ATGCTGCCMNNGCCTC |
| G4 For Lib | GAGGCGATNNKAGCATTGC |
| G4 Rev Lib | GCAATGCTMNNATCGCCTC |
| S5 For Lib | AGGCGATGGCNNKATTGCGGAAT |
| S5 Rev Lib | ATTCCGCAATMNNGCCATCGCCT |
| I6 For Lib | GATGGCAGCNNKGCGGAATAC |
| I6 Rev Lib | GTATTCCGCMNNGCTGCCATC |
| A7 For Lib | CGATGGCAGCATTNNKGAATACTTTAACC |
| A7 Rev Lib | GGTTAAAGTATTCMNNAATGCTGCCATCG |
| Y9 For Lib | GCATTGCGGAANNKTTTAACCGTCC |
| Y9 Rev Lib | GGACGGTTAAAMNNTTCCGCAATGC |
| F10 For Lib | GAATACNNKAACCGTCCGA |
| F10 Rev Lib | TCGGACGGTTMNNGTATTC |
| N11 For Lib | CGGAATACTTTNNKCGTCCGATGCA |
| N11 Rev Lib | TGCATCGGACGMNNAAAGTATTCCG |
| R12 For Lib | GGAATACTTTAACNNKCCGATGCATATTC |
| R12 Rev Lib | GAATATGCATCGGMNNGTTAAAGTATTCC |
| P13 For Lib | GGAATACTTTAACCGTNNKATGCATATTCATGATT |
| P13 Rev Lib | AATCATGAATATGCATMNNACGGTTAAAGTATTCC |
| M14 For Lib | TTTAACCGTCCGNNKCATATTCATGA |
| M14 Rev Lib | TCATGAATATGMNNCGGACGGTTAAA |
| H15 For Lib | ACCGTCCGATGNNKATTCATGAT |
| H15 Rev Lib | ATCATGAATMNNCATCGGACGGT |
| I16 For Lib | GTCCGATGCATNNKCATGATTGGCA |
| I16 Rev Lib | TGCCAATCATGMNNATGCATCGGAC |
| H17 For Lib | CGATGCATATTNNKGATTGGCAGAT |
| H17 Rev Lib | ATCTGCCAATCMNNAATATGCATCG |
| D18 For Lib | GATGCATATTCATNNKTGGCAGATTATGG |
| D18 Rev Lib | CCATAATCTGCCAMNNATGAATATGCATC |
| W19 For Lib | ATGCATATTCATGATNNKCAGATTATGGATAGC |
| W19 Rev Lib | GCTATCCATAATCTGMNNATCATGAATATGCAT |
| Q20 For Lib | ATATTCATGATTGGNNKATTATGGATAGCGG |
| Q20 Rev Lib | CCGCTATCCATAATMNNCCAATCATGAATAT |
| I21 For Lib | GATTGGCAGNNKATGGATAGCG |

| | |
|---|---|
| I21 Rev Lib | CGCTATCCATMNNCTGCCAATC |
| M22 For Lib | TGGCAGATTNNKGATAGCGGC |
| M22 Rev Lib | GCCGCTATCMNNAATCTGCCA |
| D23 For Lib | GGCAGATTATGNNKAGCGGCTATTA |
| D23 Rev Lib | TAATAGCCGCTMNNCATAATCTGCC |
| S24 For Lib | CAGATTATGGATNNKGGCTATTATGGC |
| S24 Rev Lib | GCCATAATAGCCMNNATCCATAATCTG |
| G25 For Lib | AGATTATGGATAGCNNKTATTATGGCTGAAA |
| G25 Rev Lib | TTTCAGCCATAATAMNNGCTATCCATAATCT |
| Y26 For Lib | GGATAGCGGCNNKTATGGCTGAA |
| Y26 Rev Lib | TTCAGCCATAMNNGCCGCTATCC |
| Y27 For Lib | ATAGCGGCTATNNKGGCTGAAAGCT |
| Y27 Rev Lib | AGCTTTCAGCCMNNATAGCCGCTAT |
| G28 For Lib | TAGCGGCTATTATNNKTGAAAGCTTAATT |
| G28 Rev Lib | AATTAAGCTTTCAMNNATAATAGCCGCTA |
| pAT8 G2F Lib For | CAATGGGATTTGATGGCAGC |
| pAT8 G2F Lib Rev | GCTGCCATCAAATCCCATTG |
| pAT8 G2I Lib For | CAATGGGAATTGATGGCAGC |
| pAT8 G2I Lib Rev | GCTGCCATCAATTCCCATTG |
| pAT8 G2M Lib For | CAATGGGAATGGATGGCAGC |
| pAT8 G2M Lib Rev | GCTGCCATCCATTCCCATTG |
| pAT8 G2N Lib For | CAATGGGAAATGATGGCAGC |
| pAT8 G2N Lib Rev | GCTGCCATCATTTCCCATTG |
| pAT8 G2W Lib For | CAATGGGATGGGATGGCAGC |
| pAT8 G2W Lib Rev | GCTGCCATCCCATCCCATTG |
| pAT8 G2Y Lib For | CAATGGGATATGATGGCAGC |
| pAT8 G2Y Lib Rev | GCTGCCATCATATCCCATTG |
| pAT8 A7P For | ATGGCAGCATTCCGGAATACTTTAA |
| pAT8 A7P Rev | TTAAAGTATTCCGGAATGCTGCCAT |
| pAT8 N11W For | GGAATACTTTTGGCGTCCGATGC |
| pAT8 N11W Rev | GCATCGGACGCCAAAAGTATTCC |
| pAT8 R12F For | GGAATACTTTAACTTTCCGATGCATATTC |
| pAT8 R12F Rev | GAATATGCATCGGAAAGTTAAAGTATTCC |
| pAT8 M14N For | CTTTAACCGTCCGAATCATATTCATGA |
| pAT8 M14N Rev | TCATGAATATGATTCGGACGGTTAAAG |
| pAT8 I16D For | GTCCGATGCATGATCATGATTGG |
| pAT8 I16D Rev | CCAATCATGATCATGCATCGGAC |
| pAT8 I16E For | GTCCGATGCATGAGCATGATTGG |
| pAT8 I16E Rev | CCAATCATGCTCATGCATCGGAC |
| pAT8 H17G For | CGATGCATATTGGTGATTGGCAGA |
| pAT8 H17G Rev | TCTGCCAATCACCAATATGCATCG |
| A7G N11M For | CATTGGTGAATACTTTATGCGTCCGATG |
| A7G N11M Rev | CATCGGACGCATAAAGTATTCACCAATG |

| | |
|---|---|
| S5H I16P For v2 | GATGGCCATATTGCGGAATACTTTAACCGTCCGATGCATCCGCA TGATTG |
| S5H I16P Rev v2 | CAATCATGCGGATGCATCGGACGGTTAAAGTATTCCGCAATATG GCCATC |
| N11W H17T For | CGATGCATATTACCGATTGGCAGAT |
| N11W H17T Rev | ATCTGCCAATCGGTAATATGCATCG |
| A7P I16A For | CGATGCATGCGCATGATTGGCA |
| A7P I16A Rev | TGCCAATCATGCGCATGCATCG |
| R12F W19G For | CATATTCATGATGGTCAGATTATGGATAGCG |
| R12F W19G Rev | CGCTATCCATAATCTGACCATCATGAATATG |
| S5T I16D For | GATGGCACCATTGCGGA |
| S5T I16D Rev | TCCGCAATGGTGCCATC |
| I16E D23A For | CAGATTATGGCGAGCGGCTATT |
| I16E D23A Rev | AATAGCCGCTCGCCATAATCTG |
| R12V H17G For | GGAATACTTTAACGTGCCGATGCATATT |
| R12V H17G Rev | AATATGCATCGGCACGTTAAAGTATTCC |
| pAT8 G4S For | AGGCGATAGCAGCATT |
| pAT8 G4S Rev | AATGCTGCTATCGCCT |
| pAT8 N11P For | GAATACTTTCCGCGTCCGATGCA |
| pAT8 N11P Rev | TGCATCGGACGCGGAAAGTATTC |
| pAT8 D23R For | GCAGATTATGCGTAGCGGCTATT |
| pAT8 D23R Rev | AATAGCCGCTACGCATAATCTGC |
| pAT8 D23L For | GCAGATTATGCTGAGCGGCTATT |
| pAT8 D23L Rev | AATAGCCGCTCAGCATAATCTGC |
| pAT8 D23S For | GCAGATTATGAGCAGCGGCTATT |
| pAT8 D23S Rev | AATAGCCGCTGCTCATAATCTGC |
| pAT8 I16C For | CGATGCATTGCCATGATTGGC |
| pAT8 I16C Rev | GCCAATCATGGCAATGCATCG |
| pQE-80 *Eco*RI For | GTGAGCGGATAACAATTTCACACAGAATTCATTAAAGAGG |
| pQE-80 *Hin*dIII Rev | GGATCTATCAACAGGAGTCCAAGCTCAGCTAATTAAG |
| P5-AT1F | AATGATACGGCGACCACCGAGATCTACACCGTGCGACAATGGG A |
| P7-AT1R1 | CAAGCAGAAGACGGCATACGAGATCGTGATGCTCAGCTAATTAA GCTTTCA |
| P7-AT1R2 | CAAGCAGAAGACGGCATACGAGATACATCGGCTCAGCTAATTAA GCTTTCA |
| P7-AT1R3 | CAAGCAGAAGACGGCATACGAGATGCCTAAGCTCAGCTAATTAA GCTTTCA |
| P7-AT1R4 | CAAGCAGAAGACGGCATACGAGATTGGTCAGCTCAGCTAATTAA GCTTTCA |
| P7-AT1R5 | CAAGCAGAAGACGGCATACGAGATCACTGTGCTCAGCTAATTAA GCTTTCA |
| P7-AT1R6 | CAAGCAGAAGACGGCATACGAGATATTGGCGCTCAGCTAATTAA GCTTTCA |
| P7-AT1R7 | CAAGCAGAAGACGGCATACGAGATGATCTGGCTCAGCTAATTAA GCTTTCA |

| | |
|---|---|
| P7-AT1R8 | CAAGCAGAAGACGGCATACGAGATTCAAGTGCTCAGCTAATTAAGCTTTCA |
| P7-AT1R9 | CAAGCAGAAGACGGCATACGAGATCTGATCGCTCAGCTAATTAAGCTTTCA |
| P7-AT1R10 | CAAGCAGAAGACGGCATACGAGATAAGCTAGCTCAGCTAATTAAGCTTTCA |
| P7-AT1R11 | CAAGCAGAAGACGGCATACGAGATGTAGCCGCTCAGCTAATTAAGCTTTCA |
| P7-AT1R12 | CAAGCAGAAGACGGCATACGAGATTACAAGCTCAGCTAATTAAGCTTTCA |
| P7-AT1R13 | CAAGCAGAAGACGGCATACGAGATTTGACTGCTCAGCTAATTAAGCTTTCA |
| P7-AT1R14 | CAAGCAGAAGACGGCATACGAGATGGAACTGCTCAGCTAATTAAGCTTTCA |
| P7-AT1R15 | CAAGCAGAAGACGGCATACGAGATTGACATGCTCAGCTAATTAAGCTTTCA |
| P7-AT1R16 | CAAGCAGAAGACGGCATACGAGATGGACGGGCTCAGCTAATTAAGCTTTCA |

**Table S7.** Plasmids used in this study.

| Plasmid Name | Template for PCR | Mutagenic Forward Primer for Cloning | Mutagenic Reverse Primer for Cloning | Vector backbone |
|---|---|---|---|---|
| pWC99 | N/A | N/A | N/A | N/A |
| pUboABC | pWC99 | N/A | N/A | pWC99 |
| pUboBC | N/A | N/A | N/A | pQE-80 |
| pUboABC G2F | pWC99 | pAT8 G2F Lib For | pAT8 G2F Lib Rev | pUboABC |
| pUboABC G2I | pWC99 | pAT8 G2I Lib For | pAT8 G2I Lib Rev | pUboABC |
| pUboABC G2M | pWC99 | pAT8 G2M Lib For | pAT8 G2M Lib Rev | pUboABC |
| pUboABC G2N | pWC99 | pAT8 G2N Lib For | pAT8 G2N Lib Rev | pUboABC |
| pUboABC G2W | pWC99 | pAT8 G2W Lib For | pAT8 G2W Lib Rev | pUboABC |
| pUboABC G2Y | pWC99 | pAT8 G2Y Lib For | pAT8 G2Y Lib Rev | pUboABC |
| pUboABC A7P | pWC99 | pAT8 A7P For | pAT8 A7P Rev | pUboABC |
| pUboABC N11W | pWC99 | pAT8 N11W For | pAT8 N11W Rev | pUboABC |
| pUboABC R12F | pWC99 | pAT8 R12F For | pAT8 R12F Rev | pUboABC |
| pUboABC M14N | pWC99 | pAT8 M14N For | pAT8 M14N Rev | pUboABC |
| pUboABC I16D | pWC99 | pAT8 I16D For | pAT8 I16D Rev | pUboABC |
| pUboABC I16E | pWC99 | pAT8 I16E For | pAT8 I16E Rev | pUboABC |
| pUboABC H17G | pWC99 | pAT8 H17G For | pAT8 H17G Rev | pUboABC |
| pUboABCD A7P | pWC99 | pAT8 A7P For | pAT8 A7P Rev | pWC99 |
| pUboABCD N11W | pWC99 | pAT8 N11W For | pAT8 N11W Rev | pWC99 |
| pUboABCD R12F | pWC99 | pAT8 R12F For | pAT8 R12F Rev | pWC99 |
| pUboABCD M14N | pWC99 | pAT8 M14N For | pAT8 M14N Rev | pWC99 |
| pUboABCD I16D | pWC99 | pAT8 I16D For | pAT8 I16D Rev | pWC99 |
| pUboABCD I16E | pWC99 | pAT8 I16E For | pAT8 I16E Rev | pWC99 |
| pUboABCD H17G | pWC99 | pAT8 H17G For | pAT8 H17G Rev | pWC99 |
| pUboABC A7G N11M | pWC99 | A7G N11M For | A7G N11M Rev | pUboABC |
| pUboABC S5H I16P | pWC99 | S5H I16P For v2 | S5H I16P Rev v2 | pUboABC |
| pUboABC A7P I16A | pUboABCD A7P | A7P I16A For | A7P I16A Rev | pUboABC |
| pUboABC N11W H17T | pUboABCD N11W | N11W H17T For | N11W H17T Rev | pUboABC |
| pUboABC R12F W19G | pUboABCD R12F | R12F W19G For | R12F W19G Rev | pUboABC |
| pUboABC S5T I16D | pUboABCD I16D | S5T I16D For | S5T I16D Rev | pUboABC |

| | | | | |
|---|---|---|---|---|
| pUboABC I16E D23A | pUboABCD I16E | I16E D23A For | I16E D23A Rev | pUboABC |
| pUboABC R12V H17G | pUboABCD H17G | R12V H17G For | R12V H17G Rev | pUboABC |
| pUboABCD A7G N11M | pWC99 | A7G N11M For | A7G N11M Rev | pWC99 |
| pUboABCD S5H I16P | pWC99 | S5H I16P For v2 | S5H I16P Rev v2 | pWC99 |
| pUboABCD A7P I16A | pUboABCD A7P | A7P I16A For | A7P I16A Rev | pWC99 |
| pUboABCD N11W H17T | pUboABCD N11W | N11W H17T For | N11W H17T Rev | pWC99 |
| pUboABCD R12F W19G | pUboABCD R12F | R12F W19G For | R12F W19G Rev | pWC99 |
| pUboABCD S5T I16D | pUboABCD I16D | S5T I16D For | S5T I16D Rev | pWC99 |
| pUboABCD I16E D23A | pUboABCD I16E | I16E D23A For | I16E D23A Rev | pWC99 |
| pUboABCD R12V H17G | pUboABCD H17G | R12V H17G For | R12V H17G Rev | pWC99 |
| pUboABC G4S | pWC99 | pAT8 G4S For | pAT8 G4S Rev | pUboABC |
| pUboABC N11P | pWC99 | pAT8 N11P For | pAT8 N11P Rev | pUboABC |
| pUboABC D23R | pWC99 | pAT8 D23R For | pAT8 D23R Rev | pUboABC |
| pUboABC G4S D23L | pUboABC G4S | pAT8 D23L For | pAT8 D23L Rev | pUboABC |
| pUboABC N11P D23S | pUboABC N11P | pAT8 D23S For | pAT8 D23S Rev | pUboABC |
| pUboABC I16C D23R | pUboABC D23R | pAT8 I16C For | pAT8 I16C Rev | pUboABC |

**Table S8.** Mutagenic primers used for constructing residue-specific libraries.

| Residue-Specific Library | Mutagenic Forward Primer | Mutagenic Reverse Primer |
|---|---|---|
| G2 Single Mutant | G2 For Lib | G2 Rev Lib |
| D3 Single Mutant | D3 For Lib | D3 Rev Lib |
| G4 Single Mutant | G4 For Lib | G4 Rev Lib |
| S5 Single Mutant | S5 For Lib | S5 Rev Lib |
| I6 Single Mutant | I6 For Lib | I6 Rev Lib |
| A7 Single Mutant | A7 For Lib | A7 Rev Lib |
| Y9 Single Mutant | Y9 For Lib | Y9 Rev Lib |
| F10 Single Mutant | F10 For Lib | F10 Rev Lib |
| N11 Single Mutant | N11 For Lib | N11 Rev Lib |
| R12 Single Mutant | R12 For Lib | R12 Rev Lib |
| P13 Single Mutant | P13 For Lib | P13 Rev Lib |
| M14 Single Mutant | M14 For Lib | M14 Rev Lib |
| H15 Single Mutant | H15 For Lib | H15 Rev Lib |
| I16 Single Mutant | I16 For Lib | I16 Rev Lib |
| H17 Single Mutant | H17 For Lib | H17 Rev Lib |
| D18 Single Mutant | D18 For Lib | D18 Rev Lib |
| W19 Single Mutant | W19 For Lib | W19 Rev Lib |
| Q20 Single Mutant | Q20 For Lib | Q20 Rev Lib |
| I21 Single Mutant | I21 For Lib | I21 Rev Lib |
| M22 Single Mutant | M22 For Lib | M22 Rev Lib |
| D23 Single Mutant | D23 For Lib | D23 Rev Lib |
| S24 Single Mutant | S24 For Lib | S24 Rev Lib |
| G25 Single Mutant | G25 For Lib | G25 Rev Lib |
| Y26 Single Mutant | Y26 For Lib | Y26 Rev Lib |
| Y27 Single Mutant | Y27 For Lib | Y27 Rev Lib |
| G28 Single Mutant | G28 For Lib | G28 Rev Lib |
| D3 Double Mutant | D3 For Lib | D3 Rev Lib |
| S5 Double Mutant | S5 For Lib | S5 Rev Lib |
| N11 Double Mutant | N11 For Lib | N11 Rev Lib |
| R12 Double Mutant | R12 For Lib | R12 Rev Lib |
| P13 Double Mutant | P13 For Lib | P13 Rev Lib |
| M14 Double Mutant | M14 For Lib | M14 Rev Lib |
| H15 Double Mutant | H15 For Lib | H15 Rev Lib |
| I16 Double Mutant | I16 For Lib | I16 Rev Lib |
| H17 Double Mutant | H17 For Lib | H17 Rev Lib |
| D18 Double Mutant | D18 For Lib | D18 Rev Lib |
| W19 Double Mutant | W19 For Lib | W19 Rev Lib |
| Q20 Double Mutant | Q20 For Lib | Q20 Rev Lib |
| I21 Double Mutant | I21 For Lib | I21 Rev Lib |

| | | |
|---|---|---|
| M22 Double Mutant | M22 For Lib | M22 Rev Lib |
| D23 Double Mutant | D23 For Lib | D23 Rev Lib |

**Table S9**. HPLC collection windows.

| Ubonodin Variant | HPLC Round 1 Collection Window | HPLC Round 2 Collection Window | Solvent for Dissolving Pure Peptide |
|---|---|---|---|
| A7P | 15.32 min-15.55 min | 10.65 min – 11.2 min | water |
| R12F | 17.15 min-17.7 min | N/A | 50/50 water/acetonitrile |
| M14N | 13.95 min – 14.2 min | 7.5 min - 7.8 min | water |
| I16D | 13.8 min – 14.2 min | 7. 45 min – 7.9 min | water |
| I16E | 13.8 min - 14.15 min | 7.55 min - 8 min | water |
| H17G | 15.8 min – 16.1 min | N/A | 50/50 water/acetonitrile |
| A7G N11M | 15.8 min – 16.05 min | N/A | 50/50 water/acetonitrile |
| S5H I16P | 13.6 min – 14 min | 21.5 min – 21.8 min | water |
| A7P I16A | 14.15 min-14.8 min | N/A | water |
| R12F W19G | 15.9 min-16.5 min | 13.15 min -14.8 min | 50/50 water/acetonitrile |
| S5T I16D | 13.9 min – 14.3 min | 7.6 min – 8 min | water |
| I16E D23A | 13.8 min – 14.3 min | 7.2 min – 7.8 min | water |
| R12V H17G | 17.4 min -17.8 min | N/A | 50/50 water/acetonitrile |
